# Parallel Arg/N-degron recognition systems regulate coenzyme A biosynthesis

**DOI:** 10.64898/2026.09.15.751905

**Authors:** Xiaolu Wang, Boju Wang, Liang Zhang, Shuxin Yan, Nixian Zhao, Yueling Zhao, Fengying Shi, Shiqi Yin, Dongxing Chen, Ying Yu, Cheng Dong, Wenyi Mi

**Affiliations:** Key Laboratory of Breast Cancer Prevention and Therapy (Ministry of Education), The Province and Ministry Co-sponsored Collaborative Innovation Center for Medical Epigenetics, Key Laboratory of Immune Microenvironment and Disease (Ministry of Education), School of Basic Medical Sciences, Tianjin Medical University, Tianjin 300070, China; Department of Pharmacology, Tianjin Key Laboratory of Inflammatory Biology, Center for Cardiovascular Diseases, Tianjin Medical University, Tianjin 300070, China; Department of Medicinal Chemistry, Tianjin Key Laboratory on Technologies Enabling Development of Clinical Therapeutics and Diagnostics, School of Pharmacy, Tianjin Medical University, Tianjin, China

## Abstract

Coenzyme A (CoA) is an essential metabolic cofactor whose biosynthesis is controlled by pantothenate kinases (PANKs), the rate-limiting enzymes in CoA synthesis. Although CoA production is extensively regulated, whether PANK abundance is controlled by selective protein degradation remains unknown. Here, we identify PANK1β and PANK3 as substrates of the Arg/N-degron pathway. Two mechanistically distinct E3 ligase systems, UBR4-KCMF1 and MKLN1-CTLH, independently converge on a common N-terminal MK Arg/N-degron to promote PANK degradation and restrict CoA biosynthesis. UBR4-KCMF1 recognizes the MK Arg/N-degron through KCMF1 ZZ domain, whereas CTLH complex employs the B30.2-domain proteins RanBP9 and RanBP10 as previously unrecognized Arg/N-degron recognins. Structural, biochemical, and cellular analyses demonstrate that RanBP9 and RanBP10 recognize both Met-X and canonical Arg/N-degrons through their B30.2 domains, using a mechanism distinct from canonical UBR proteins. Together, these findings establish a new branch of the mammalian Arg/N-degron pathway and reveal ubiquitin-dependent control of PANK abundance in CoA homeostasis.

## Introduction

The central role of coenzyme A (CoA) in cellular metabolism has been recognized since its discovery more than 70 years ago as the cofactor responsible for carrying “activated acetate”.^1^ CoA lies at the intersection of catabolism, anabolism, and metabolic signaling, serving as an essential cofactor for numerous biological processes, including the tricarboxylic acid (TCA) cycle, nutrient oxidation, histone acetylation, and the biosynthesis of lipids, glycans, and heme.^2^

De novo CoA biosynthesis is highly conserved across all domains of life and proceeds through a nearly universal five-step enzymatic pathway. In the first and rate-limiting step, pantothenate (vitamin B5) is phosphorylated by pantothenate kinases (PANKs) to generate 4′-phosphopantothenate.^3^ The human genome encodes three catalytically active PANK paralogs, PANK1-3. Two active PANK1 isoforms, PANK1α and PANK1β, are generated through alternative splicing and translation initiation.^4–6^ These enzymes exhibit distinct subcellular localizations, with PANK1β and PANK3 residing predominantly in the cytosol, PANK1α in the nucleus, and PANK2 in the mitochondrial intermembrane space.^7^

As with many metabolic pathways, CoA biosynthesis is regulated at multiple levels. These mechanisms include subcellular compartmentalization,^8^ feedback inhibition by CoA thioesters and related metabolites,^9–13^ transcriptional and translational regulation,^14–17^ and post-translational modifications such as phosphorylation.^2,18^ Together, these mechanisms dynamically coordinate CoA production with metabolic demands and nutrient availability. Whether PANK abundance is itself controlled through selective protein degradation, however, remains unknown. Given the widespread role of ubiquitin-mediated proteolysis in regulating enzyme abundance and metabolic flux,^19,20^ regulated degradation of PANKs may represent an additional mechanism governing CoA homeostasis.

The ubiquitin-proteosome system (UPS) is the principal pathway for selective protein degradation in eukaryotic cells.^21–25^ Within the UPS, E3 ubiquitin ligases confer substrate specificity by catalyzing ubiquitin conjugation to target proteins, most commonly through lysine-linked polyubiquitin chains.^26,27^ Substrate recognition is dictated by degradation signals, or degrons, intrinsic sequence or structural elements that direct protein turnover.^28^ Among the earliest degrons to be characterized were N-degrons, destabilizing N-terminal residues that trigger rapid proteasomal degradation.^29,30^ N-degron pathways comprise several distinct branches that recognize different classes of destabilizing N termini, including the Arg/N-degron, Ac/N-degron, Pro/N-degron, fMet/N-degron, and GASTC/N-degron pathways.^23,31^

In 2017, Chen and colleagues identified the *S. cerevisiae* GID4 protein, a subunit of the GID ubiquitin ligase complex, as the N-recognin of the Pro/N-degron pathway.^19^ The mammalian counterpart of the yeast GID ubiquitin ligase, the CTLH (C-terminal to LisH) E3 ligase complex, has since been implicated in erythropoiesis, embryogenesis, microtubule dynamics, chromosome segregation, and other cellular processes.^19,32–34^ Structurally, the CTLH is assembled around a catalytic MAEA–RMND5A RING heterodimer together with the scaffold proteins TWA1, RanBP9, and ARMC8.^35–38^ Incorporation of WDR26 or MKLN1 generates distinct supramolecular CTLH assemblies that contain multiple substrate-recognition modules. Among these, GID4 recognizes Pro/N-degron substrates such as HMGCS1,^20,32,39^ WDR26 functions as a substrate receptor for proteins including NMNAT1 and HBP1,^36,40,41^ and MKLN1 has been implicated in degradation of UNG2 through FAM72A and, more recently, in a C-degron pathway.^42,43^ Despite these advances, the full repertoire of CTLH substrate receptors and the degron pathways governed by this complex remain incompletely understood.

Within the Arg/N-degron pathway, members of the UBR family, including UBR1,UBR2, UBR4, and UBR5, function as canonical Arg/N-recognins.^31^ These E3 ligases recognize proteins bearing destabilizing N-terminal residues, including Arg, Lys, His, Ile, Leu, Phe, Tyr, Trp. In addition, initiator methionine followed by bulky hydrophobic residues (Met-Φ; Φ = Ile, Leu, Phe, Tyr, or Trp) constitutes a distinct class of Arg/N-degrons recognized by UBR proteins.^44–46^ More recently, N-terminal methionine followed by Arg or Lys was identified as a potential destabilizing motif targeted by UBR family E3 ligases,^47^ suggesting that the substrate-recognition landscape of the Arg/N-degron pathway is broader than previously appreciated.

Here, we identify PANK proteins as previously unrecognized substrates of N-degron pathway and establish regulated proteolysis as an additional layer of CoA homeostasis control. We show that two mechanistically distinct E3 ligase systems, the UBR4-KCMF1 heterodimeric ligase and the MKLN1-containing CTLH complex, independently recognize a common N-terminal MK Arg/N-degron to cooperatively regulate the stability of PANK1β and PANK3, thereby controlling de novo CoA biosynthesis. Mechanistically, we identify RanBP9 and RanBP10 as a previously unrecognized class of Arg/N-degron recognins that mediates substrate recognition by the CTLH complex through their B30.2 domains. Structural, biochemical, and cellular analyses further reveal that RanBP9 and RanBP10 recognize both Met-X and canonical Arg/N-degrons through a substrate-recognition mechanism distinct from that of canonical UBR-family proteins. Together, our findings establish the CTLH complex as a previously unrecognized branch of the mammalian Arg/N-degron pathway, expand the repertoire of Arg/N-degron recognition mechanism, and uncover a fundamental mechanism by which ubiquitin-dependent proteolysis is coupled to CoA metabolism.

## Results

### A CRISPR screen identifies two E3 ligases that control PANK stability

To determine whether PANK abundance is regulated through selective protein degradation, we established stable cell lines expressing epitope-tagged PANK1α, PANK1β, PANK2, and PANK3 (Figures 1A and 1B). Inhibition of the ubiquitin–proteasome system (UPS) with either the ubiquitin-activating enzyme (E1) inhibitor MLN7243 or the proteasome inhibitor MG132 markedly increased the abundance of the predominantly cytosolic isoforms PANK1β and PANK3, whereas inhibition of lysosomal degradation had no detectable effect. By contrast, the abundance of nuclear isoform PANK1α and mitochondrial isoform PANK2 remained largely unchanged following inhibition of either the UPS or lysosomal pathway (Figure 1B), indicating that selective ubiquitin-dependent degradation preferentially regulates the cytosolic PANK isoforms.

**Figure 1.**
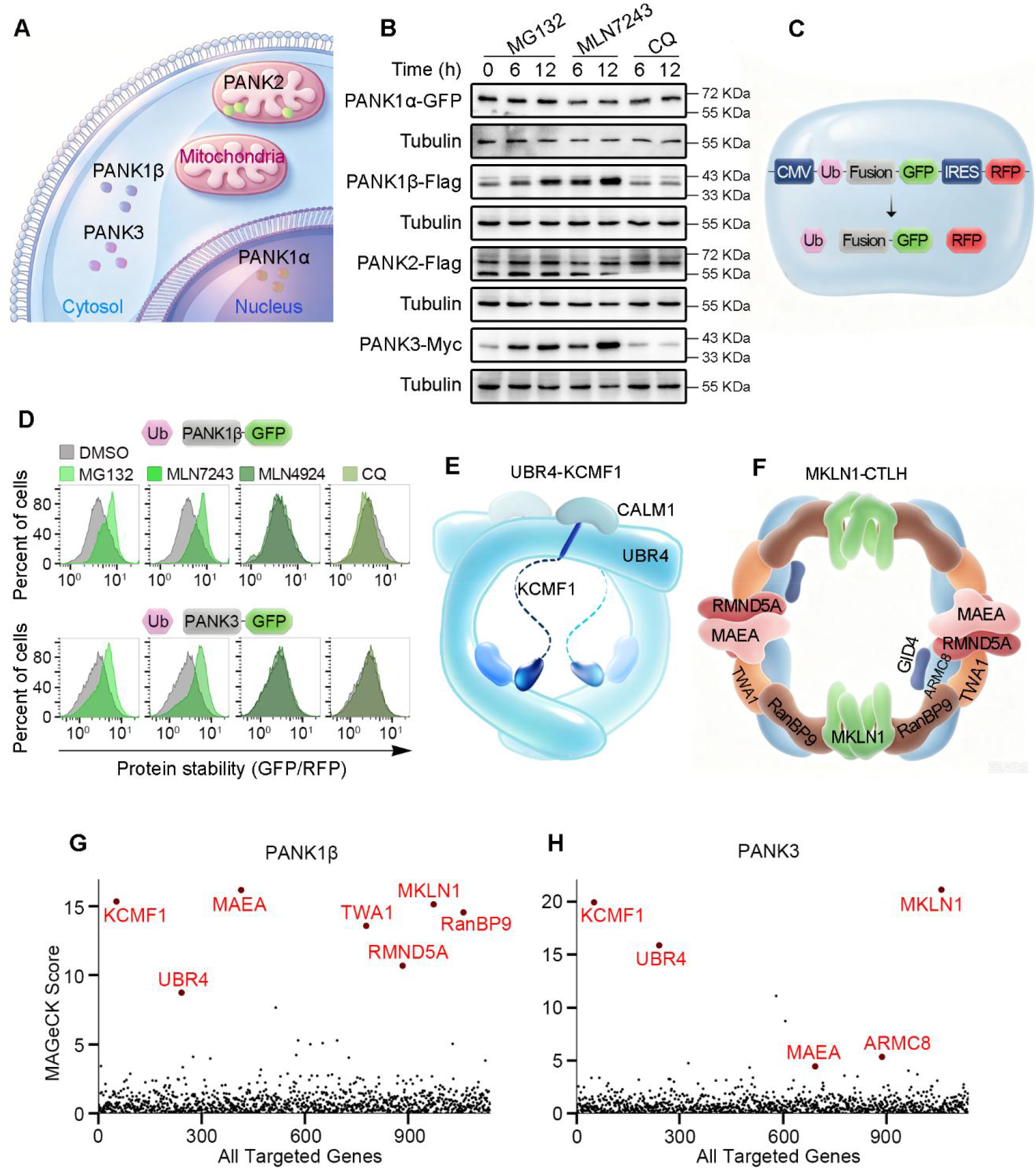
A CRISPR screen identifies the UBR4–KCMF1 and MKLN1-CTLH as candidate regulators of PANK stability. (A) Schematic illustration of the subcellular localization of human PANK paralogs (PANK1-3). (B) Stable HEK293T cell lines expressing epitope-tagged PANK proteins were treated with 10 μM MG132, MLN7243 or chloroquine (CQ) for the indicated times. Cell lysates were analyzed by immunoblotting using antibodies against epitope tags, with tubulin serving as a loading control. (C) Schematic illustration of the global protein stability (GPS) reporter system, showing the design of the construct and the resulting protein products. (D) Stable HeLa cell lines ectopically expressing GPS reporter for PANK1β or PANK3 were treated with 10μM MG132, MLN7243, MLN4924 or CQ for 6 h. Relative stability PANK proteins was quantified by flow cytometry based on the GFP/RFP fluorescence ratio. (E) Simplified schematic of the UBR4-KCMF1 complex based on published cryo-electron microscopy (cryo-EM) structure. (F) Simplified schematic of MKLN1-CTLH complex based on published cryo-EM structure. (G) Results of CRISPR screens identifying E3 ligases regulating PANK1β degradation, analyzed via MAGeCK. (H) Results of CRISPR screens identifying E3 ligases regulating PANK3 degradation, analyzed via MAGeCK.

To independently validate these observations, we employed the global protein stability (GPS) reporter system,^47^ in which individual PANK proteins were fused to GFP and co-expressed with red fluorescent protein (RFP) through an internal ribosome entry site (IRES), allowing protein stability to be quantified by the GFP/RFP fluorescence ratio (Figure 1C). Consistent with the immunoblot analyses, treatment with either MG132 or MLN7243 strongly stabilized PANK1β and PANK3, whereas PANK2 remained unaffected (Figures 1D and S1A). Together, these findings establish the ubiquitin-proteasome system (UPS) as a major regulator of PANK1β and PANK3 turnover.

To identify the E3 ligases responsible for PANK degradation, we constructed a pooled CRISPR-Cas9 library targeting 1,118 genes implicated in protein degradation, with 10 sgRNAs per gene, and performed pooled genetic screens using the GPS reporter system (Figure S1B and Table S1). Multiple components of two independent E3 ligase complexes emerged among the strongest hits. One was the UBR4-KCMF1 complex (Figures 1E, 1G, and 1H), in which KCMF1 is an essential cofactor required for UBR4 E3 ligase activity.^48,49^ The other was the CTLH complex (Figures 1F, 1G, and 1H). In the PANK1β screen, multiple CTLH subunits representing distinct functional modules were robustly enriched, including the catalytic components MAEA and RMND5A, the scaffold proteins TWA1 and RanBP9, and the supramolecular assembly factor MKLN1.^33,37^ Although fewer CTLH components were recovered in the PANK3 screen, both screens independently identified the UBR4–KCMF1 and CTLH complexes as candidate regulators of PANK stability. The recurrent enrichment of multiple subunits from these two E3 ligase systems strongly suggested that UBR4–KCMF1 and the MKLN1-containing CTLH complex are previously unrecognized regulators of PANK turnover, providing the foundation for the mechanistic studies described below.

### UBR4–KCMF1 recognizes an N-terminal MK motif as an Arg/N-degron

To validate the screening results, we genetically disrupted components of the UBR4-KCMF1 complex using CRISPR-Cas9. Depletion of either UBR4 or KCMF1 markedly increased the stability of PANK1β and PANK3 in the GPS reporter assay (Figures S2A and S2B), and this phenotype was recapitulated in independent clonal knockout cell lines (Figures 2A and 2B). Conversely, re-expression of KCMF1 in KCMF1-deficient cells restored PANK1β degradation (Figure 2C), confirming the requirement of the UBR4-KCMF1 complex for efficient PANK turnover.

**Figure 2.**
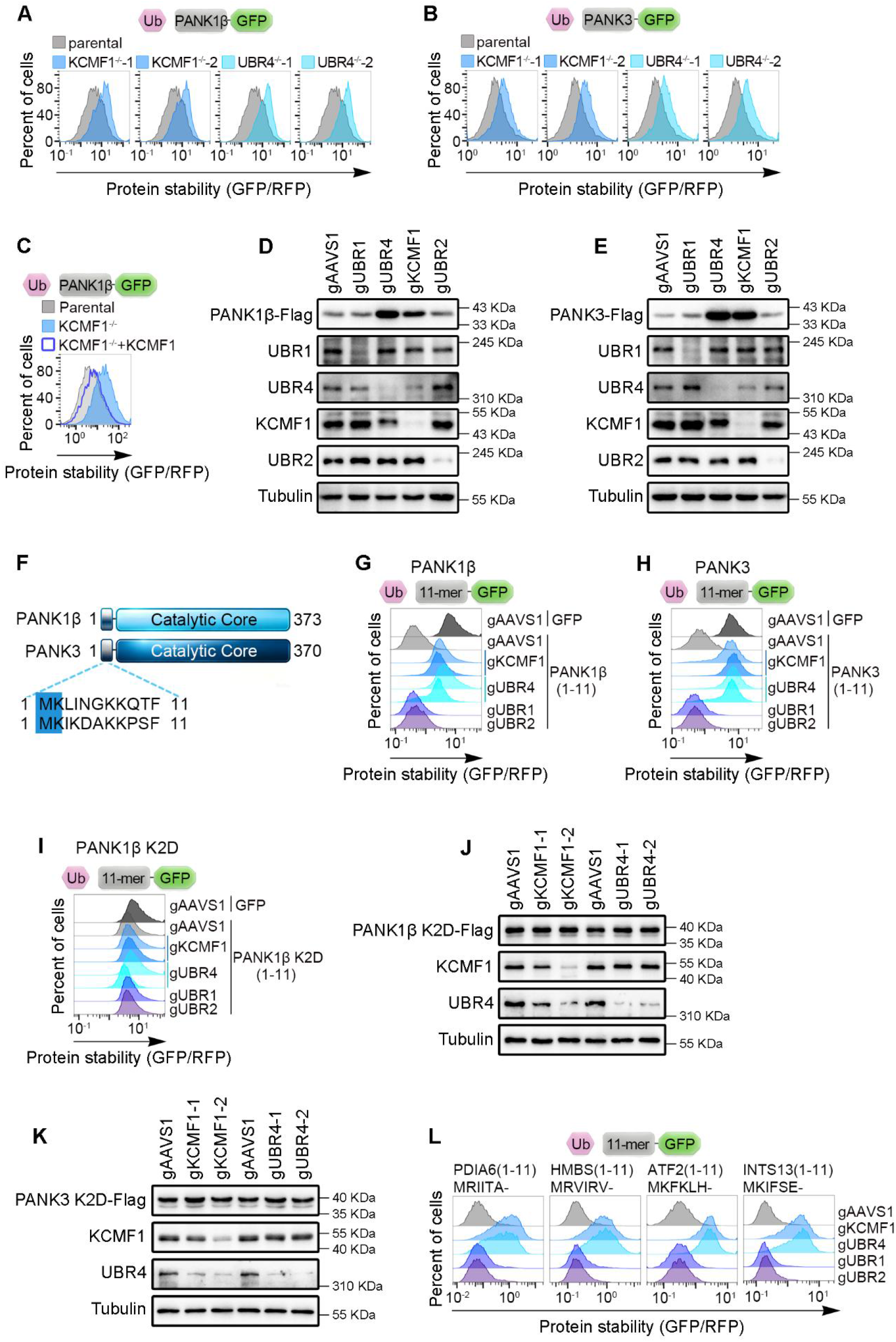
UBR4–KCMF1 recognizes an N-terminal MK motif as an Arg/N-degron. (A) WT, KCMF1^-/-^ and UBR4^-/-^ clonal HEK293T cells were transfected with the PANK1β-GFP, as indicated, and analyzed by flow cytometry. Two independent knockout clones were examined for each genotype. (B) WT, KCMF1^-/-^ and UBR4^-/-^ clonal HEK293T cells were transfected with the PANK3-GFP reporter, as indicated, and analyzed by flow cytometry. Two independent knockout clones were examined for each genotype. (C) WT and KCMF1^-/-^ clonal HEK293T cells, with or without re-expression of KCMF1, were transfected with the PANK1β-GFP reporter and analyzed by flow cytometry. (D) Stable HEK293T cells ectopically expressing PANK1β-Flag were transduced with control gRNA or gRNA targeting UBR1, UBR4, KCMF1 or UBR2, as indicated. Cell lysates were analyzed by immunoblotting with the indicated antibodies. (E) Stable HEK293T cells ectopically expressing PANK3-Flag were transduced with control gRNA or gRNA targeting UBR1, UBR4, KCMF1 or UBR2, as indicated. Cell lysates were analyzed by immunoblotting with the indicated antibodies. (F) Schematic illustration of human PANK1β and PANK3 proteins structure, showing the alignment of their N-terminal 11 amino acids sequences. The two conserved N-terminal residues shared by PANK1β and PANK3 are highlighted in blue. Numbers indicate the total length of each protein. (G) Stable HEK293T cells transduced with control gRNA or gRNAs targeting KCMF1, UBR4, UBR1 or UBR2 were transfected with the GFP alone or PANK1β(1-11aa)-GFP reporter, as indicated, and analyzed by flow cytometry. Two independent gRNAs were used for KCMF1and UBR4. (H) Stable HEK293T cells transduced with control gRNA or gRNA targeting KCMF1, UBR4, UBR1 or UBR2 were transfected with the GFP alone or PANK3(1-11aa)-GFP reporter, as indicated, and analyzed by flow cytometry. Two independent gRNAs were used for KCMF1and UBR4. (I) Stable HEK293T cells transduced with control gRNA or gRNA targeting KCMF1, UBR4, UBR1 or UBR2 were transfected with the GFP alone or PANK1β K2D(1-11aa)-GFP reporter, as indicated, and analyzed by flow cytometry. Two independent gRNAs were used for KCMF1and UBR4. (J) Stable HEK293T cell lines ectopically expressing Flag-tagged full-length PANK1β K2D protein were transduced with control gRNA or gRNA targeting KCMF1 or UBR4, as indicated. Cell lysates were analyzed by immunoblotting with the indicated antibodies. (K) Stable HEK293T cell lines ectopically expressing Flag-tagged full-length PANK3 K2D protein were transduced with control gRNA or gRNA targeting KCMF1 or UBR4, as indicated. Cell lysates were analyzed by immunoblotting with the indicated antibodies. (L) Stable HEK293T cells transduced with control gRNA or gRNA targeting KCMF1, UBR4, UBR1 or UBR2 were transfected with GFP reporters bearing the indicated N-terminal MK- or MR-initiating peptides and analyzed by flow cytometry.

Immunoblot analysis of PANK isoforms fused to a small Flag epitope yielded similar results. Depletion of either UBR4 or KCMF1 substantially increased the abundance of PANK1β and PANK3, whereas depletion of UBR1 or UBR2 had no detectable effect (Figures 2D and 2E). In contrast, the abundance of nuclear PANK1α and mitochondrial PANK2 remained largely unchanged following UBR4 or KCMF1 depletion (Figures S2C and S2D). These findings establish UBR4-KCMF1 as a selective regulator of the stability of the predominantly cytosolic PANK1β and PANK3 isoforms.

We next sought to define the determinant that mediates recognition of PANK proteins by UBR4–KCMF1. Notably, N-terminal fusion of GFP to either PANK1β or PANK3 abolished their responsiveness to UBR4 or KCMF1 depletion (Figures S2E, S2F, and S2G), suggesting that the native N termini of PANK1β and PANK3 contain a degradation signal required for UBR4–KCMF1-dependent turnover. Consistent with this possibility, fusion of the first 11 amino acids of either PANK1β or PANK3 to GFP markedly destabilized the reporter compared with GFP alone (Figures 2F, 2G, and 2H). This destabilization was abolished by loss of UBR4 or KCMF1 but was unaffected by depletion of UBR1 or UBR2 (Figures 2G and 2H), indicating that the N terminal sequences of PANK1β and PANK3 are sufficient to confer UBR4-KCMF1-dependent degradation.

Both PANK1β and PANK3 initiate with methionine followed by lysine (MK) (Figure 2F). Given our finding that their N termini are sufficient to confer UBR4-KCMF1-dependent degradation, together with recent evidence identifying MK- and MR-initiating sequences as potential substrates of UBR-family E3 ligases,^47^ we hypothesized that the N-terminal MK motif constitutes the degron recognized by UBR4–KCMF1. To test this possibility, we substituted the second-position lysine with the negatively charged residue aspartate (K2D). The K2D substitution markedly impaired degradation of the N-terminal GFP reporter and abolished its stabilization in response to UBR4 or KCMF1 depletion (Figure 2I). Similarly, Flag-tagged PANK1β (K2D) and PANK3 (K2D) no longer accumulated following loss of UBR4 or KCMF1 (Figures 2J and 2K). Furthermore, multiple independent GFP reporters bearing representative MK- or MR-initiating N-terminal peptides were stabilized upon loss of UBR4 or KCMF1 (Figure 2L). Taken together, these findings identify N-terminal methionine followed by lysine or arginine as a class of Arg/N-degrons targeted by the UBR4–KCMF1 E3 ligase complex and establish the PANK N terminus as the MK Arg/N-degron that directs UBR4–KCMF1-dependent turnover.

### The KCMF1 ZZ domain mediates recognition of the destabilizing MK Arg/N-degron

Recent cryo-EM studies have shown that the UBR4-KCMF1 complex forms a ∼1.3-MDa ring-shaped assembly containing a central substrate-binding cavity.^44,48–50^ Within this cavity, two substrate recognition modules—the UBR domain of UBR4 and the ZZ domain of KCMF1—have been implicated in the recognition of N-terminal Arg/N-degrons (Figure 3A). To determine how UBR4-KCMF1 recognizes the N-terminal MK Arg/N-degron of PANK1β and PANK3, we first examined the contribution of the UBR domain. Surprisingly, genetic excision of the UBR domain in UBR4 had no detectable effect on PANK1β stability, despite the marked stabilization observed following complete loss of UBR4 (Figures 3B and S3A). These findings indicate that the UBR domain of UBR4 is dispensable for recognition of the MK degron of PANK1β.

**Figure 3.**
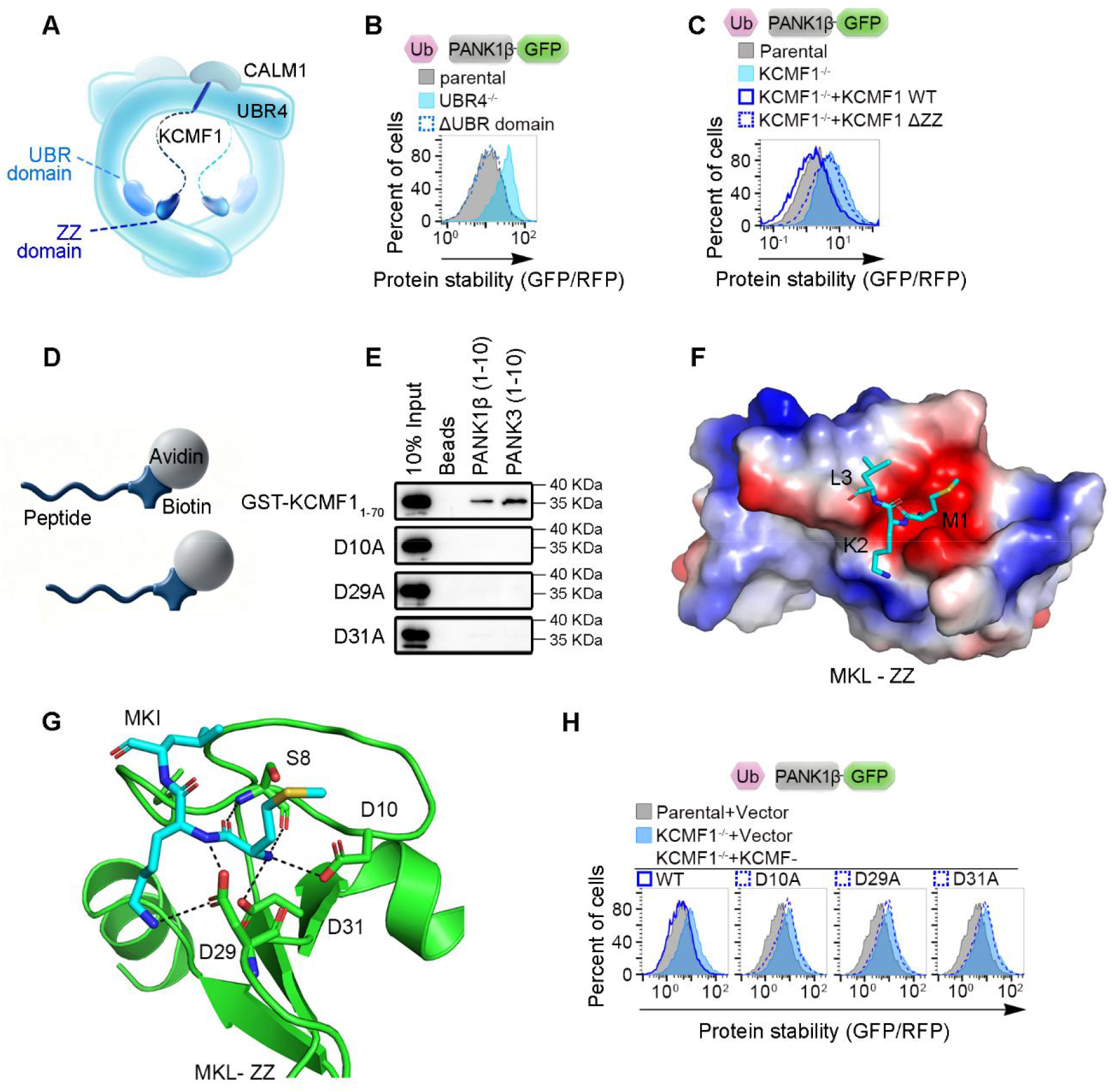
The KCMF1 ZZ domain mediates recognition of the destabilizing MK Arg/N-degron. (A) Schematic illustration of the UBR4-KCMF1 complex, highlighting the UBR domain of UBR4, and the ZZ domain of KCMF1. (B) WT, UBR4^-/-^, UBR4 UBR domain-excised clonal HEK293T cells were transfected with the PANK1β-GFP reporter, as indicated, and analyzed by flow cytometry. (C) WT, KCMF1^-/-^ and KCMF1^-/-^ HEK293T cells reconstituted with WT KCMF1 or a ZZ domain deletion mutant (ΔZZ) as indicated, were transfected with the PANK1β-GFP reporter and analyzed by flow cytometry. (D) Schematic illustration of biotinylated peptides. (E) Peptide pull-down using indicated biotinylated N-terminal peptides of PANK1β and PANK3, incubated with GST-fused WT or mutant KCMF1 ZZ domain. (F) The AlphaFold3-predicted electrostatic potential surface of KCMF1 ZZ domain in complex with N-terminal MKL motif of PANK1β. Positive and negative electrostatic potentials are shown in blue and red, respectively; the MKL peptide is shown as turquoise sticks. (G) The AlphaFold3-predicted detailed interactions between the KCMF1 ZZ domain and the PANK1β N-terminal MKL peptide. The MKL peptide is shown as turquoise sticks, and KCMF1 residues are colored green. (H) WT, KCMF1^-/-^ and KCMF1^-/-^ HEK293T cells reconstituted with WT KCMF1 or the indicated ZZ domain point mutants were transfected with the PANK1β-GFP reporter and analyzed by flow cytometry.

We next investigated whether the KCMF1 ZZ domain mediates recognition of the MK degron. Re-expression of wild-type KCMF1 in KCMF1-deficient cells fully restored PANK1β degradation, whereas re-expression of a ZZ-domain deletion mutant failed to do so (Figures 3C and S3B), demonstrating that the KCMF1 ZZ domain is essential for UBR4-KCMF1-dependent degradation of PANK1β.

To determine whether the ZZ domain directly recognizes the MK degron, we performed peptide pull-down assays using bacterially purified GST-tagged KCMF1 ZZ domain and biotinylated N-terminal peptides derived from PANK1β and PANK3. The ZZ domain efficiently bound both MK-containing peptides in vitro (Figures 3D and 3E), supporting a direct interaction between the KCMF1 ZZ domain and the N-terminal MK degron. To gain structural insight into this interaction, we generated an AlphaFold 3 model of the KCMF1 ZZ domain in complex with the PANK1β N-terminal peptide (Figure 3F). The model positioned the MK degron within a negatively charged surface pocket of the ZZ domain, resembling the experimentally determined structure of the ZZ domain bound to a canonical Arg/N-degron peptide.^48^

The predicted interface highlighted several conserved acidic residues within the ZZ domain that could contribute to recognition of the positively charged MK degron (Figure 3G). Consistent with this model, alanine substitution of key acidic residues, D10, D29, or D31, markedly impaired binding of the ZZ domain to the MK-containing peptides in vitro (Figure 3E). Moreover, whereas wild-type KCMF1 restored PANK1β degradation in KCMF1-deficient cells, the corresponding ZZ-domain mutants failed to rescue degradation, as measured by the Ub-GPS reporter assay (Figures 3H and S3C). Together, these biochemical, structural, and structure-guided genetic analyses establish the KCMF1 ZZ domain as the substrate-recognition module for the N-terminal MK Arg/N-degron and reveal a mechanism by which UBR4-KCMF1 targets PANK proteins for ubiquitin-dependent degradation.

### The MKLN1-CTLH complex promotes PANK degradation through MK Arg/N-degron recognition

To validate the screening results, we systematically disrupted components of the supramolecular donut-shaped CTLH complex using CRISPR-Cas9. Consistent with the screen, depletion of the catalytic module components MAEA and RMND5A, as well as the scaffold proteins TWA1 and RanBP9, markedly stabilized PANK1β in the GPS reporter assay (Figure 4A). In contrast, disruption of the ARMC8-GID4 module, which functions as the Pro/N-degron recognin within the CTLH complex, had no detectable effect on PANK1β stability. Notably, loss of MKLN1, one of the two supramolecular assembly factors of the CTLH complex, strongly stabilized PANK1β, whereas depletion of the other assembly factor, WDR26, had no effect (Figure 4A). PANK3, in contrast, was only modestly stabilized upon depletion of core CTLH components, including MAEA, TWA1, or RanBP9 (Figure S4A). Given the robust effects observed for PANK1β, we focused subsequent mechanistic studies on this isoform.

**Figure 4.**
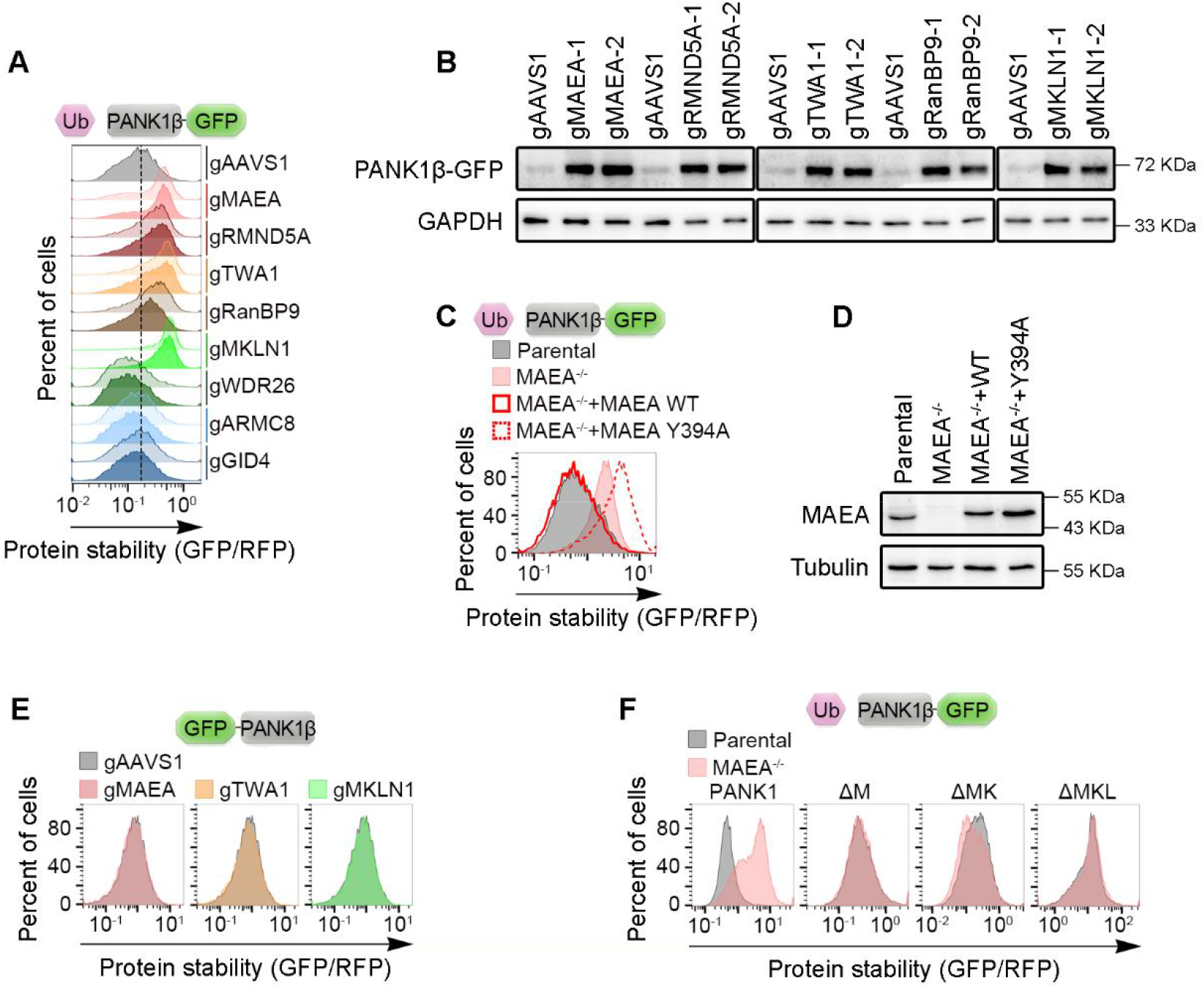
The MKLN1-CTLH complex promotes PANK degradation through MK Arg/N-degron recognition. (A) WT and knockout Hela cells lacking the indicated individual CTLH complex components were transfected with PANK1β-GFP reporter and analyzed by flow cytometry. Two independent gRNAs were used to target each gene. (B) Stable HeLa cells ectopically expressing PANK1β-GFP were transduced with control gRNA or gRNA targeting MAEA, RMND5A, TWA1, RanBP9, or MKLN1, as indicated. Two independent gRNAs were used to target each gene. Cell lysates were analyzed by immunoblotting with the indicated antibodies. (C) WT and MAEA^-/-^ HEK293T cells, together with MAEA^-/-^ cells reconstituted with WT MAEA or the Y394A mutant, were transfected with the PANK1β-GFP reporter and analyzed by flow cytometry. (D) Cell lysates from WT, MAEA^-/-^, and MAEA^-/-^ HEK293T cells re-expressing either WT MAEA or Y394A mutant were analyzed by immunoblotting with the indicated antibodies. (E) Stable HeLa cells transduced with control gRNA or gRNA targeting MAEA, TWA1 or MKLN1, as indicated, were transfected with GFP-PANK1β reporter, and reporter stability was assessed by flow cytometry. (F) WT and MAEA−/− clonal HeLa cells were transfected with the indicated PANK1β-GFP reporters, including the WT reporter and variants lacking the N-terminal M, MK, or MKL residues, and analyzed by flow cytometry.

Immunoblot analysis of GFP-tagged PANK1β independently confirmed the reporter-based findings. Depletion of MAEA, RMND5A, TWA1, RanBP9, or MKLN1 substantially increased PANK1β abundance, whereas disruption of ARMC8, GID4, or WDR26 had little or no detectable effect (Figures 4B and S4B). To further validate these observations, we generated independent clonal knockout cell lines lacking individual CTLH subunits. Consistent with the pooled CRISPR analyses, PANK1β was dramatically stabilized in MAEA-, RanBP9-, and MKLN1-deficient cells, but remained largely unchanged in WDR26-, ARMC8-, and GID4-deficient cells (Figure S4C). We next asked whether CTLH catalytic activity is required for PANK1β degradation. Re-expression of wild-type MAEA in MAEA-deficient cells fully restored PANK1β degradation, whereas a catalytically impaired MAEA(Y394A) mutant carrying a substitution within the non-RING priming element failed to rescue degradation (Figures 4C and 4D). These results indicate that CTLH-dependent ubiquitin ligase activity is required for efficient PANK1β turnover. Collectively, these results identify the MKLN1-CTLH assembly as a major regulator of PANK1β degradation. The requirement for core CTLH catalytic and scaffold components, together with the dispensability of ARMC8 and GID4, indicates that PANK1β is targeted by a CTLH substrate-recognition mechanism distinct from the canonical Pro/N-degron pathway.

We next sought to define the substrate determinant recognized by MKLN1-CTLH complex. N-terminal fusion of GFP to PANK1β completely abolished its stabilization in response to depletion of MAEA, TWA1, or MKLN1 (Figure 4E), indicating that an accessible native N terminus is required for CTLH-dependent recognition. Consistent with this finding, deletion of the initiator methionine, the N-terminal MK dipeptide, or the MKL tripeptide each abolished CTLH-dependent degradation of PANK1β (Figure 4F). Thus, the extreme N terminus of PANK1β is both necessary for recognition and degradation by the MKLN1-CTLH complex. Notably, the requirement for the N-terminal MK motif mirrors that observed for UBR4-KCMF1-mediated PANK1β degradation, raising the possibility that these two mechanistically distinct E3 ligase systems converge on the same MK Arg/N-degron.

### RanBP9 and RanBP10 function as substrate receptors of the CTLH complex

Within the supramolecular MKLN1-CTLH assembly, GID4 is the best-characterized substrate receptor, whereas MKLN1 has also been implicated in substrate recognition.^42,43^ However, disruption of the GID4-ARMC8 module had no detectable effect on PANK1β stability, whereas loss of MKLN1 robustly stabilized PANK1β (Figure 4A). Consistent with the genetic results, pharmacological inhibition of GID4 with PFI-7, a selective GID4 inhibitor, failed to stabilize PANK1β (Figure S5A), indicating that PANK1β degradation proceeds independently of the canonical GID4-mediated recognition pathway. The selective requirement for MKLN1 therefore raised the possibility whether MKLN1 functions as the substrate receptor recruiting PANK1β to the CTLH complex.

To test this possibility, we examined the association of PANK1β with the CTLH complex by co-immunoprecipitation. As expected, PANK1β associated with the core CTLH subunits MAEA and RanBP9 in control cells, and these interactions remained readily detectable in either GID4- or ARMC8-deficient cells (Figure S5B), further supporting the dispensability of the canonical GID4-ARMC8 recognition module. Unexpectedly, loss of MKLN1 did not disrupt the association of PANK1β with either MAEA or RanBP9 (Figure 5A), indicating that MKLN1 is not required for recruitment of PANK1β to the CTLH complex. Furthermore, PANK1β retained associated with RanBP9 in MAEA-deficient cells (Figure 5B), suggesting that RanBP9 itself may engage PANK1β independently of the catalytic module and thereby contribute directly to substrate recruitment.

**Figure 5.**
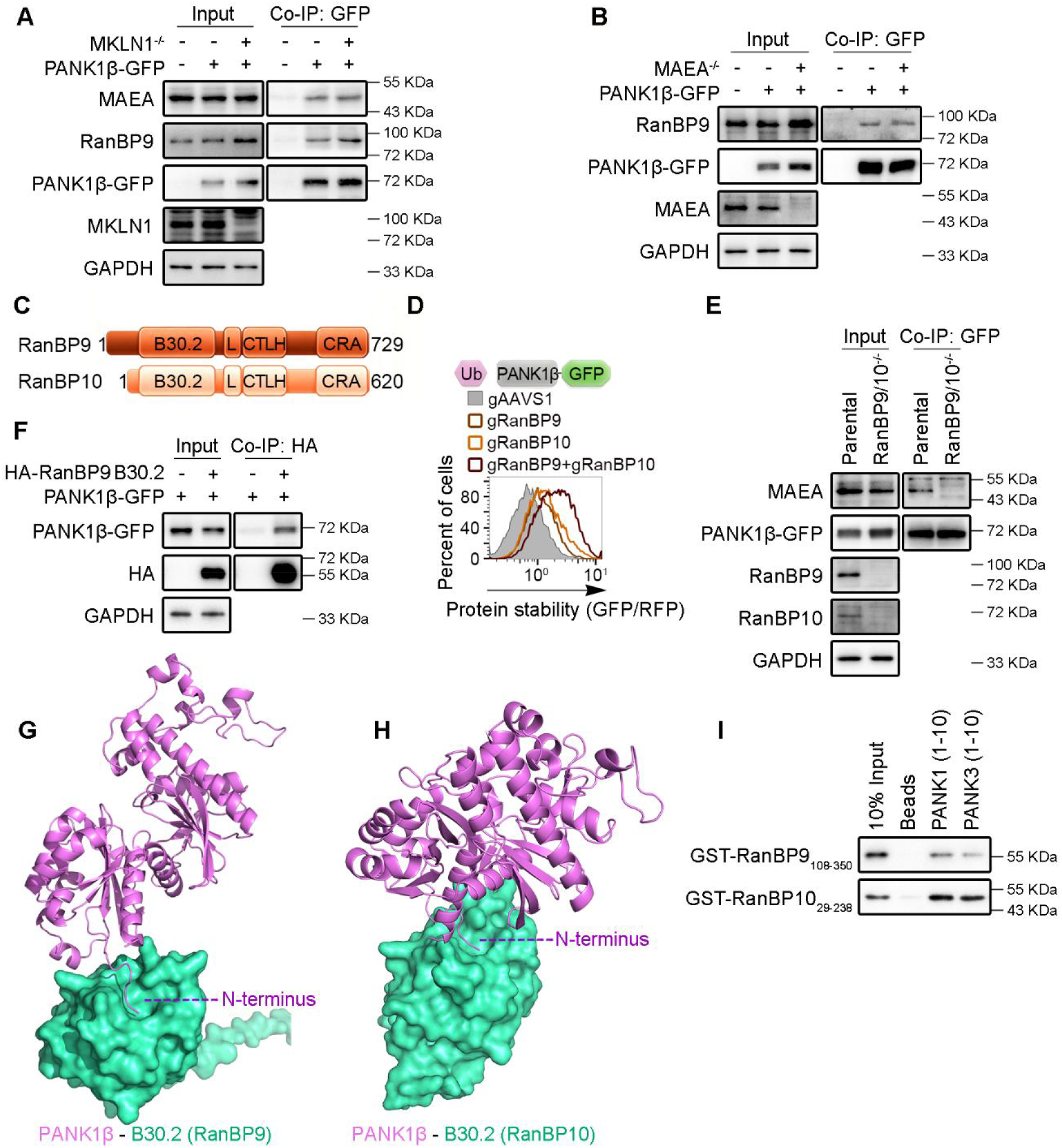
RanBP9 and RanBP10 function as substrate receptors of the CTLH complex. (A) Parental HeLa and MKLN1^-/-^ clonal cells transiently expressing vector control or PANK1β-GFP were subjected to anti-GFP immunoprecipitation, followed by immunoblotting for endogenous MAEA and RanBP9. Input and immunoprecipitated samples were analyzed with the indicated antibodies. (B) HEK293T parental or MAEA^-/-^ clonal cells transiently expressing vector or PANK1β-GFP were subjected to anti-GFP co-IP to examine interactions with endogenous RanBP9. Input and co-IP samples were analyzed by immunoblotting with the indicated antibodies. (C) Schematic illustration of the domain architecture of human RanBP9 and RanBP10, including the B30.2, LisH (L), CTLH and CRA domains. Numbers indicate the total length of each protein. (D) Stable HeLa cells transduced with control gRNA or gRNA targeting RanBP9, RanBP10 or both genes simultaneously, as indicated, were transfected with PANK1β-GFP reporter and analyzed by flow cytometry. (E) HEK293T parental or RanBP9-/-RanBP10-/- clonal cells transiently expressing PANK1β-GFP were subjected to anti-GFP co-IP to examine interactions with endogenous MAEA. Input and co-IP samples were analyzed by immunoblotting with the indicated antibodies. (F) HEK293T cells transiently co-expressing an empty vector or HA-tagged RanBP9 B30.2 domain (residues 1-359) together with PANK1β-GFP were subjected to anti-HA co-IP. Input and co-IP samples were analyzed by immunoblotting with the indicated antibodies. (G) Alphafold3-predicted structure of the RanBP9 B30.2 domain in complex with full-length PANK1β. The PANK1β is shown in purple, with its N-terminus labeled, and the B30.2 domain is colored green. (H) Alphafold3-predicted structure of the RanBP10 B30.2 domain in complex with full-length PANK1β. The PANK1β is shown in purple, with its N-terminus labeled, and the B30.2 domain is colored green. (I) Peptide pull-down assay using indicated biotinylated N-terminal peptides of PANK1β and PANK3, incubated with GST-fused RanBP9 or RanBP10 B30.2 domain. Figure 6

RanBP9 and RanBP10 are closely related paralogs that arose from an ancestral gene duplication event and share highly conserved domain architectures (Figure 5C). Structural and biochemical studies have shown that these paralogs occupy equivalent positions within the CTLH assemblies.^51,52^ We therefore asked whether RanBP10, like RanBP9, contributes to PANK1β degradation. Knockout of either RanBP9 or RanBP10 resulted in partial stabilization of PANK1β, whereas simultaneous depletion of both proteins produced substantially greater stabilization (Figure 5D), indicating that RanBP9 and RanBP10 function redundantly to promote CTLH-dependent PANK1β turnover. Consistently, simultaneous deletion of RanBP9 and RanBP10 abolished the association of PANK1β with the catalytic subunit MAEA (Figure 5E), suggesting that these paralogs mediate recruitment of PANK1β to the CTLH complex.

Within the CTLH assembly, RanBP9 and RanBP10 serve as central organizational scaffolds. Their LisH-CRA^C^ domains recruit TWA1 to form RanBP9/10-TWA1 heterodimers, whereas the CTLH-CRA^N^ domain mediates incorporation of MKLN1 into the complex.^37,38^ Notably, structural models place the N-terminal B30.2 domain of RanBP9 and RanBP10 toward the central cavity of the donut-shaped CTLH assembly, exposing a prominent interaction surface whose function remains poorly understood. This architecture, together with the genetic and interaction data above, promoted us to hypothesize that the B30.2 domain serves as a substrate-recognition module for PANK1β.

To test this possibility, we expressed an isolated HA-tagged B30.2 domain and examined its association with PANK1β by co-immunoprecipitation. Similar to full-length RanBP9, the isolated B30.2 domain efficiently associated with PANK1β (Figure 5F), indicating that the B30.2 domain is sufficient for substrate binding. Strikingly, N-terminal fusion of GFP to PANK1β completely abolished its interaction with the B30.2 domain (Figure S5C), suggesting that the B30.2 domain recognizes a determinant within the native PANK1β N terminus. Consistent with this possibility, AlphaFold 3^53^ predicted a direct interaction between the intrinsically disordered N-terminal region of PANK1β and the B30.2 domains of RanBP9 and RanBP10 (Figures 5G and 5H). We next tested this interaction biochemically using peptide pull-down assays with biotinylated N-terminal peptides derived from PANK1β and PANK3 and bacterially purified GST-tagged B30.2 domains. Both RanBP9 and RanBP10 B30.2 domains bound directly to the N-terminal peptides of PANK1β and PANK3 in vitro, with a stronger interaction observed for the PANK1β peptide (Figure 5I). Collectively, these findings identify the B30.2 domains of RanBP9 and RanBP10 as substrate-recognition modules that directly engage the N-terminal MK Arg/N-degron of PANK proteins, thereby providing a molecular explanation for how the complex targets PANK proteins for degradation.

### Structural basis of RanBP9 recognition of the MK Arg/N-degron

To elucidate the structural basis of MK Arg/N-degron recognition, we determined the crystal structure of RanBP9 in complex with the N-terminal segment of PANK1β (Figures 6A and S6A). The RanBP9 B30.2 domain adopts a canonical β-sandwich architecture composed of two antiparallel β-sheets flanked by N- and C-terminal helices. A prominent ligand-binding pocket is formed by surface-exposed loops on one face of the β-sandwich (Figure S6A). The N-terminal MKL segment of PANK1β is accommodated within this pocket through an extensive network of hydrogen-bonding interactions (Figure 6B). Notably, the free α-amino group of the initiator methionine (M1) is anchored within the pocket through hydrogen bonds with D257, D258, and H255, whereas the methionine carbonyl oxygen is coordinated by the indole nitrogen of W247. Moreover, the side chain of M1 forms van der Waals interactions with Tyr229. Although K2 does not form direct hydrogen bonds with the protein, the binding pocket is highly acidic, which may favor accommodation of the positively charged K2 side chain. The backbone amide of the third residue, L3, additionally forms a hydrogen bond with the carbonyl oxygen of G266. Additionally, L3 forms hydrophobic interaction with the W247. Together, these interactions create a complementary binding interface that engages both the free N terminus and the adjacent residues of PANK1β, thereby securing the MK Arg/N-degron within the B30.2 pocket.

**Figure 6.**
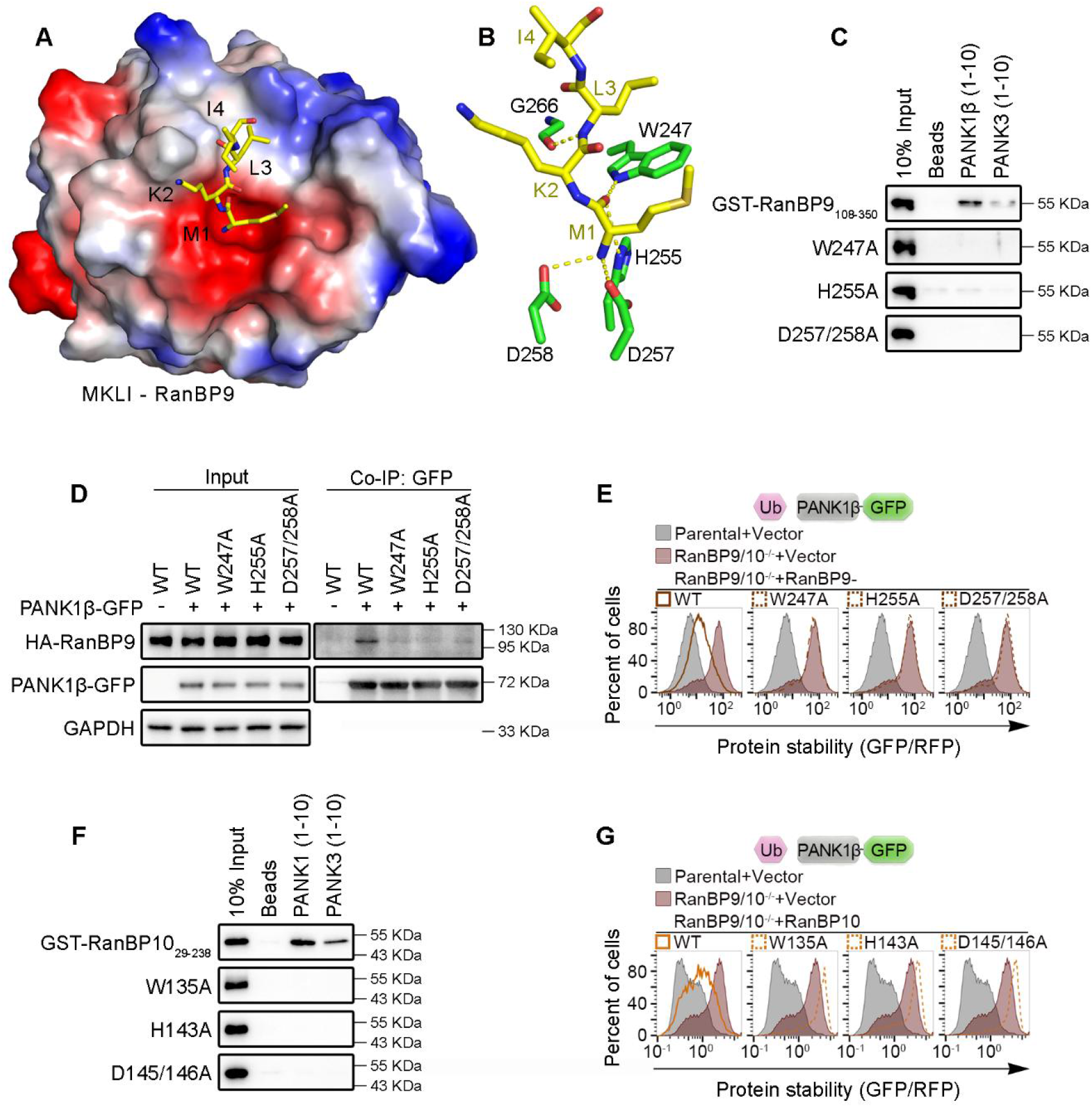
Structural basis of RanBP9 recognition of the MK Arg/N-degron. (A) Electrostatic potential surface of RanBP9 B30.2 domain in complex with N-terminal MKLI motif of PANK1β. Positive and negative electrostatic potentials are shown in blue and red, respectively; the MKLI peptide is shown as yellow sticks. (B) Detailed interactions between the RanBP9 B30.2 domain and the PANK1β N-terminal MKLI peptide. The MKLI peptide is shown as yellow sticks, and RanBP9 residues involved in direct contacts are shown as green sticks. (C) Peptide pull-down assay using indicated biotinylated N-terminal peptides of PANK1β and PANK3, incubated with GST-fused WT or mutant RanBP9 B30.2 domains. (D) HEK293T cells transiently co-expressing HA-tagged WT or B30.2 domain mutant RanBP9 together with either an empty vector or PANK1β-GFP, as indicated, were subjected to anti-GFP co-IP. Input and co-IP samples were analyzed by immunoblotting with the indicated antibodies. (E) WT, RanBP9^-/-^RanBP10^-/-^, and RanBP9^-/-^RanBP10^-/-^ HeLa cells reconstituted with WT RanBP9 or the indicated B30.2 domain mutants were transfected with the PANK1β-GFP reporter and analyzed by flow cytometry. (F) Peptide pull-down assay using indicated biotinylated N-terminal peptides of PANK1β and PANK3, incubated with GST-fused WT or mutant RanBP10 B30.2 domains. (G) WT, RanBP9^-/-^RanBP10^-/-^, and RanBP9^-/-^RanBP10^-/-^ HeLa cells reconstituted with WT RanBP10 or the indicated B30.2 domain mutants were transfected with the PANK1β-GFP reporter and analyzed by flow cytometry.

We next sought to determine whether the residues forming this pocket are required for degron recognition and substrate degradation. *In vitro* peptide pull-down assays showed that alanine substitution of W247, H255, or D257/D258 markedly impaired or abolished binding of the RanBP9 B30.2 domain to biotinylated N-terminal peptides derived from PANK1β and PANK3 (Figure 6C). Consistent with these findings, full-length RanBP9 carrying the corresponding mutations failed to co-immunoprecipitate with PANK1β in cells (Figure 6D), identifying W247, H255, D257, and D258 as critical determinants of substrate recognition. Importantly, re-expression of wild-type RanBP9 in RanBP9/RanBP10-deficient cells restored PANK1β degradation, whereas the W247A, H255A, and D257A/D258A mutants failed to rescue degradation (Figures 6E and S6B). These results establish that the B30.2-mediated recognition interface is required not only for peptide binding but also for CTLH-dependent turnover of PANK1β.

The residues forming the degron-binding pocket are highly conserved in the paralog RanBP10 (Figure S6C). Consistent with a conserved recognition mechanism, the corresponding RanBP10 mutants—W135A, H143A, and D145A/D146A—markedly impaired binding to the N-terminal MK peptides in *in vitro* pull-down assays (Figure 6F), disrupted the interaction with PANK1β in cells (Figure S6D), and failed to restore PANK1β degradation in RanBP9/RanBP10-deficient cells (Figures 6G and S6E). Thus, RanBP9 and RanBP10 use conserved B30.2-domain pocket to recognize the N-terminal MK Arg/N-degron and promote CTLH-dependent substrate degradation. Taken together, these structural, biochemical, and genetic analyses identify RanBP9 and RanBP10 as previously unrecognized MK Arg/N-degron recognins. The crystal structure reveals how the B30.2 domain directly engages the free N-terminal MK degron, while structure-guided mutagenesis demonstrates that this interaction is essential for CTLH-mediated degradation of PANK proteins.

### UBR4–KCMF1 and MKLN1–CTLH function additively to control PANK turnover and CoA biosynthesis

The above findings identify UBR4–KCMF1 and MKLN1–CTLH as two mechanistically distinct E3 ligase complexes that converge on the same N-terminal MK Arg/N-degron to regulate PANK stability. This convergence raised the question of whether the two ligases act through a common regulatory mechanism or instead operate as independent degradation systems. We first examined whether either complex influences the abundance or integrity of the other. Disruption of the CTLH complex by deletion of either MAEA or MKLN1 had no detectable effect on the abundance of UBR4 or KCMF1 (Figure S7A). Conversely, loss of either UBR4 or KCMF1 did not alter the abundance of CTLH complex subunits (Figure S7B). These results indicate that UBR4-KCMF1 and MKLN1-CTLH are assembled and maintained independently, arguing against a hierarchical relationship between the two E3 ligase systems.

We next asked whether these two ligases function within a common degradation pathway or independently target the same MK Arg/N-degron. To distinguish these possibilities, we generated cells simultaneously lacking KCMF1 and MKLN1, thereby disrupting both UBR4–KCMF1 and MKLN1–CTLH activities (Figure 7A). Re-expression of KCMF1 restored UBR4–KCMF1 activity and was sufficient to promote PANK1β degradation despite the continued absence of MKLN1 and the resulting loss of the CTLH activity (Figure 7B). Conversely, re-expression of MKLN1 restored CTLH-dependent degradation of PANK1β despite the continued absence of KCMF1 and the resulting loss of UBR4–KCMF1 activity (Figure 7C). Thus, either E3 ligase complex was independently sufficient to promote PANK1β degradation in the absence of the other, demonstrating that UBR4–KCMF1 and MKLN1–CTLH do not function in an obligate linear pathway but instead constitute parallel degradation systems that converge on the same N-terminal MK Arg/N-degron.

**Figure 7.**
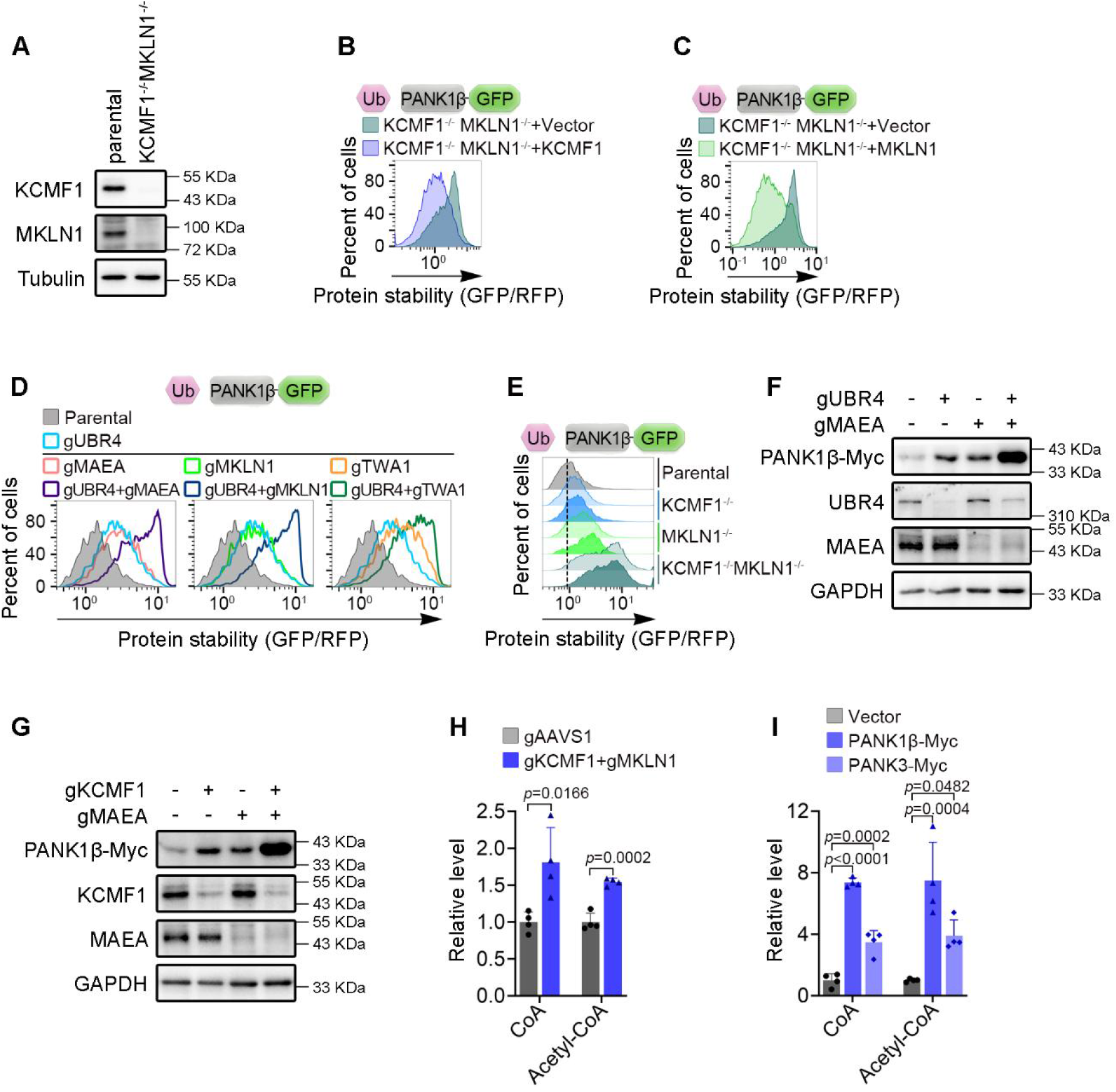
UBR4–KCMF1 and MKLN1–CTLH function additively to control PANK turnover and CoA biosynthesis. (A) Cell lysates from WT and KCMF1^-/-^MKLN1^-/-^ HeLa cells were analyzed by immunoblotting with the indicated antibodies. (B) KCMF1^-/-^MKLN1^-/-^ clonal HeLa cells and KCMF1^-/-^MKLN1^-/-^ cells reconstituted with KCMF1, as indicated, were transfected with the PANK1β-GFP reporter and analyzed by flow cytometry. (C) KCMF1^-/-^MKLN1^-/-^ clonal HeLa cells and KCMF1^-/-^MKLN1^-/-^ cells reconstituted with MKLN1, as indicated, were transfected with the PANK1β-GFP reporter and analyzed by flow cytometry. (D) HeLa cells transduced with control gRNA or gRNA targeting UBR4, individual CTLH complex components (MAEA, MKLN1 or TWA1), or the indicated combinations, were transfected with the PANK1β-GFP reporter and analyzed by flow cytometry. (E) WT, KCMF1^-/-^, MKLN1^-/-^, and KCMF1^-/-^MKLN1^-/-^ clonal HeLa cells were transfected with the PANK1β-GFP reporter and analyzed by flow cytometry. Two independent knockout clones were examined for each genotype. (F) Stable HeLa cells ectopically expressing PANK1β-Myc were transduced with control gRNA or gRNA targeting UBR4, MAEA or both genes simultaneously, as indicated. Cell lysates were analyzed by immunoblotting with the indicated antibodies. (G) Stable HeLa cells ectopically expressing PANK1β-Myc were transduced with control gRNA or gRNA targeting KCMF1, MAEA or both genes simultaneously, as indicated. Cell lysates were analyzed by immunoblotting with the indicated antibodies. (H) Metabolite levels in HEK293T cells transduced with control gRNA or gRNAs targeting both KCMF1 and MKLN1 as indicated, were quantified by LC-MS/MS and normalized to cell number. (I) Metabolite levels in HEK293T cells transduced with vector, PANK1β-Myc or PANK3-Myc as indicated, were quantified by LC–MS/MS and normalized to cell number.

We next determined whether the two E3 ligases make additive contributions to PANK turnover. Disruption of UBR4–KCMF1 through deletion of UBR4, or disruption of the CTLH complex through deletion of MAEA, MKLN1, or TWA1, resulted in partial stabilization of PANK1β. Strikingly, simultaneous disruption of both E3 ligase systems produced substantially greater stabilization than disruption of either complex alone (Figure 7D), a phenotype recapitulated in independent clonal knockout cell lines (Figure 7E). Immunoblot analysis of Myc-tagged PANK1β further confirmed these findings: loss of either E3 ligase complex partially increased PANK1β abundance, whereas combined disruption resulted in markedly greater PANK1β accumulation (Figures 7F and 7G). PANK3 exhibited a similar pattern of regulation. Disruption of either UBR4-KCMF1 or MKLN1-CTLH partially increased PANK3 abundance, whereas simultaneous disruption of both complexes produced a substantially greater increase (Figures S7C and S7D). Consistent with these findings, endogenous PANK3 abundance was similarly increased upon combined disruption of the two pathways (Figure S7E), whereas disruption of MKLN1-CTLH alone produced a more modest effect on PANK3 stability (Figures S7C-S7E). Together, these results demonstrate that UBR4–KCMF1 and MKLN1–CTLH independently contribute to PANK1β and PANK3 turnover and exert additive effects through convergent recognition of the N-terminal MK Arg/N-degron.

Because PANK1β and PANK3 catalyze the rate-limiting step of de novo CoA biosynthesis, we next asked whether the combined regulation of PANK stability by these two E3 ligases affects cellular CoA metabolism. We quantified intracellular CoA and its major derivative, acetyl-CoA, by LC–MS/MS. Simultaneous disruption of both E3 ligase complexes significantly increased the levels of both metabolites (Figures 7H and S7F), consistent with enhanced PANK abundance and activity. To determine whether elevated PANK abundance is sufficient to drive this metabolic phenotype, we measured CoA and acetyl-CoA levels in cells overexpressing PANK1β or PANK3. Overexpression of either PANK isoform similarly increased intracellular CoA and acetyl-CoA levels (Figures 7I and S7G), recapitulating the metabolic effects of simultaneous disruption of both E3 ligases. Together, these findings establish UBR4–KCMF1 and MKLN1–CTLH as parallel, additive E3 ligase systems that constrain PANK abundance and thereby limit de novo CoA biosynthesis.

### RanBP9/10 define a previously unrecognized class of Arg/N-degron recognins

N-terminal methionine followed by bulky hydrophobic residues (Ile, Leu, Phe, Tyr, or Trp) constitutes Met-Φ Arg/N-degrons recognized by UBR family E3 ligase.^44–46^ Our identification of N-terminal methionine followed by lysine or arginine as degrons recognized by UBR4-KCMF1 expands the repertoire of Met-X Arg/N-degrons (X = Ile, Leu, Phe, Tyr, Trp, Lys, or Arg). Unexpectedly, we further identified RanBP9 and RanBP10 as previously unrecognized degron receptors for the N-terminal MK motif, revealing a distinct mode of substrate recognition within the CTLH E3 ligase complex. These findings prompted us to ask whether RanBP9 and RanBP10 are selective for the MK degron or instead recognize a broader spectrum of Met-X Arg/N-degrons.

To define the substrate specificity of RanBP9 and RanBP10, we substituted the second lysine residue of PANK1β with Arg, His, Ile, Leu, Phe, Tyr, or Trp, generating MR-, MH-, MI-, ML-, MF-, MY-, and MW-starting variants. Corresponding biotinylated 10-residue N-terminal peptides were synthesized for in vitro binding assays (Figure 8A). Peptide pull-down experiments revealed that the B30.2 domains of both RanBP9 and RanBP10 bound not only the native MK peptide but also the MR-, MF-, and MW-starting peptides, whereas little or no binding was detected for the MI-, ML-, or MY-starting peptides (Figure 8B), demonstrating that RanBP9 and RanBP10 selectively recognize a subset of Met-X Arg/N-degrons. Notably, the MH-starting peptide was also efficiently recognized by the B30.2 domains of both RanBP9 and RanBP10 (Figure 8B), suggesting that their substrate-recognition specificity extends beyond the currently defined Met-X Arg/N-degrons.

**Figure 8.**
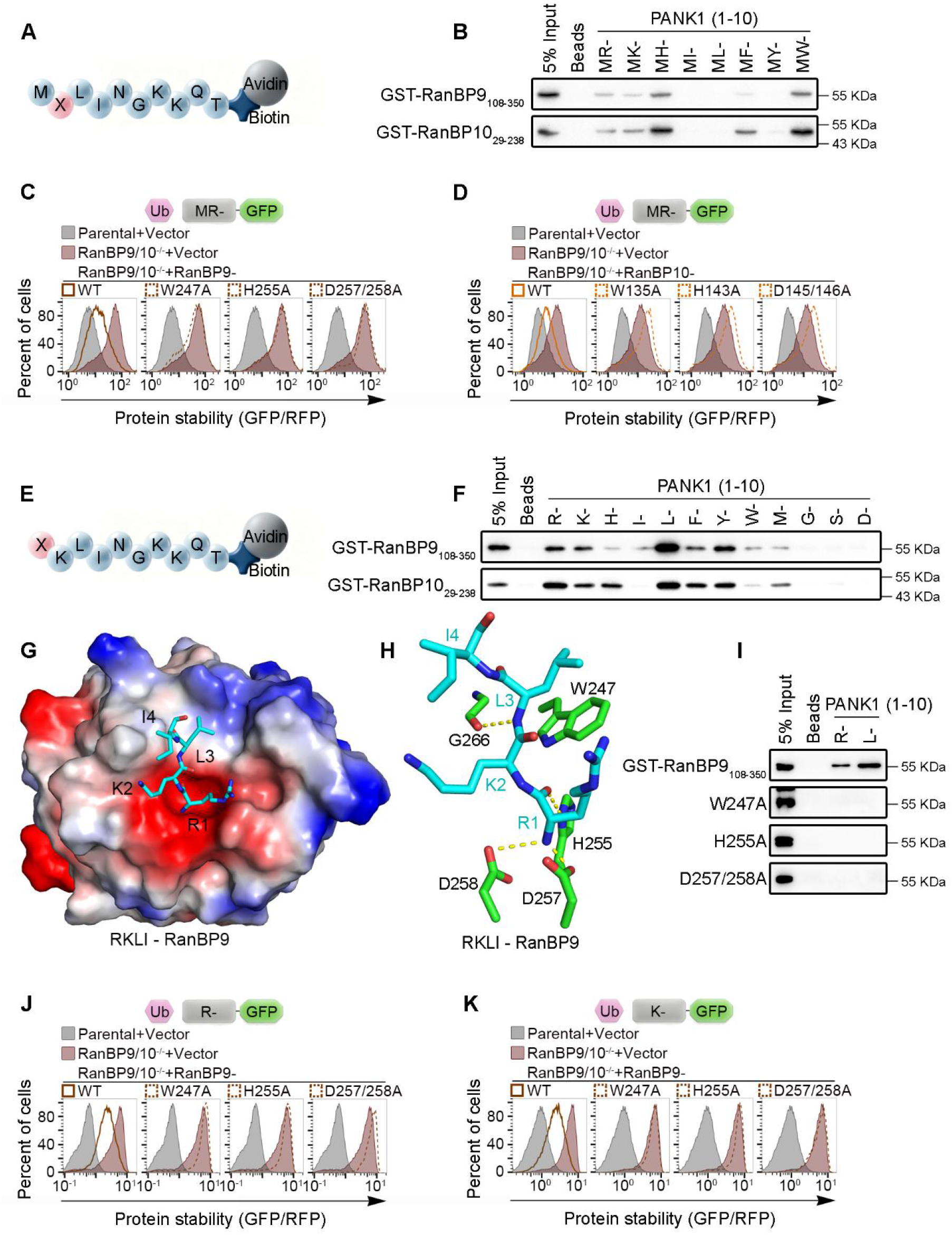
RanBP9/10 define a previously unrecognized class of Arg/N-degron recognins. (A) Schematic illustration of biotinylated peptide pull-down assay with N-terminal peptides carrying substitutions at the second residue. (B) Peptide pull-down assay using the indicated biotinylated PANK1β N-terminal peptides carrying substitutions at the second residue, incubated with GST-fused RanBP9 or RanBP10 B30.2 domains. (C) WT, RanBP9^-/-^RanBP10^-/-^, and RanBP9^-/-^RanBP10^-/-^ HeLa cells reconstituted with WT RanBP9 or the indicated B30.2 domain mutants were transfected with MR-initiating PANK1β-GFP variant and analyzed by flow cytometry. (D) WT, RanBP9^-/-^RanBP10^-/-^, and RanBP9^-/-^RanBP10^-/-^ HeLa cells reconstituted with WT RanBP10 or the indicated B30.2 domain mutants were transfected with MR-initiating PANK1β-GFP variant and analyzed by flow cytometry. (E) Schematic illustration of biotinylated peptide pull-down assay with N-terminal peptides carrying substitutions at the first residue. (F) Peptide pull-down assay using the indicated biotinylated PANK1β N-terminal peptides carrying substitutions at the first residue, incubated with GST-fused RanBP9 or RanBP10 B30.2 domains. (G) Electrostatic potential surface of RanBP9 B30.2 domain in complex with RKLI peptide. Positive and negative electrostatic potentials are shown in blue and red, respectively; the RKLI peptides are represented as turquoise sticks. (H) Detailed interactions between the RanBP9 B30.2 domain and the RKLI peptide. The RKLI peptide is shown as turquoise sticks, and RanBP9 residues involved in direct contacts are shown as green sticks. (I) Peptide pull-down using the indicated biotinylated R- and H-initiating PANK1β N-terminal peptides, incubated with GST-fused WT or mutant RanBP9 B30.2 domain. (J) WT, RanBP9^-/-^RanBP10^-/-^, and RanBP9^-/-^RanBP10^-/-^ HeLa cells reconstituted with WT RanBP9 or the indicated B30.2 domain mutants were transfected with the R-initiating PANK3-GFP variant reporter and analyzed by flow cytometry. (K) WT, RanBP9-/-RanBP10^-/-^, and RanBP9^-/-^RanBP10^-/-^ HeLa cells reconstituted with WT RanBP9 or the indicated B30.2 domain mutants were transfected with the K-starting PANK3-GFP variant reporter and analyzed by flow cytometry.

We next determined whether peptide binding correlated with degron function in cells. GPS reporter assays showed that, similar to the MK-starting PANK1β, the MR-, MH-, MF-, and MW-starting PANK1β variants were markedly stabilized upon depletion of RanBP9 and RanBP10, whereas the MY-starting variant was largely unaffected (Figure S8A). To determine whether substrate degradation depends on B30.2-mediated degron recognition, we performed complementation experiments in RanBP9/RanBP10 double-knockout cells. Re-expression of wild-type RanBP9 restored CTLH-dependent degradation of the MR- and MH-starting PANK1β variants, whereas the substrate-binding pocket mutants W247A, H255A, or D257A/D258A failed to rescue substrate degradation (Figures 8C and S8B). Likewise, wild-type RanBP10, but not the corresponding pocket mutants W135A, H143A, or D145A/D146A, restored degradation of these substrates (Figures 8D and S8C). Together, these findings demonstrate that the B30.2 domains of RanBP9 and RanBP10 directly mediate recognition of a subset of Met-X Arg/N-degrons and are required for their subsequent degradation by the CTLH E3 ligase complex.

We next asked whether RanBP9 and RanBP10 can recognize canonical Arg/N-degrons in addition to Met-X Arg/N-degrons. To this end, we replaced the initiator methionine of PANK1β with Arg, Lys, His, Ile, Leu, Phe, Tyr, or Trp, generating R-, K-, H-, I-, L-, F-, Y-, and W-starting variants (Figure 8E). Corresponding biotinylated 10-residue N-terminal peptides, together with G-, S-, and D-starting peptides as negative controls, were synthesized for in vitro binding assays. Peptide pull-down experiments revealed that the B30.2 domains of both RanBP9 and RanBP10 robustly bound R-, K-, H-, L-, F-, Y-, and W-starting peptides, whereas little or no binding was detected for the I-starting peptide or the G-, S-, or D-starting control peptides (Figure 8F). These findings demonstrate that RanBP9 and RanBP10 recognize a broad subset of canonical Arg/N-degrons. Consistent with this conclusion, introduction of representative canonical Arg/N-degrons (R, K, or L) at the N terminus of PANK3 similarly conferred binding to the B30.2 domains of both RanBP9 and RanBP10 (Figure S8D).

To elucidate the molecular basis for canonical Arg/N-degron recognition, we determined the co-crystal structure of the RanBP9 B30.2 domain in complex with an R-starting PANK1β peptide (Figure 8G). Similar to the MK degron, the N-terminal arginine is accommodated within the same surface pocket of the B30.2 domain and engages an analogous network of interactions that mediate substrate recognition (Figures 8H). Guided by the structure, we mutated key residues lining the substrate-binding pocket. In vitro pull-down assays showed that alanine substitution of W247, H255, or D257/D258 abolished binding of the RanBP9 B30.2 domain to biotinylated R- and L-starting PANK1β peptides (Figure 8I), further validating the structural model and demonstrating that the same B30.2 pocket mediates recognition of distinct Arg/N-degron classes.

We next asked whether canonical Arg/N-degron recognition by RanBP9 and RanBP10 is required for substrate degradation in cells. GPS reporter assays showed that the R-, K-, and H-starting variants of both PANK1β and PANK3 were markedly stabilized following depletion of RanBP9 and RanBP10 (Figures S8E and S8F). Complementation experiments in RanBP9/RanBP10 double-knockout cells further demonstrated that re-expression of wild-type RanBP9 restored CTLH-dependent degradation of the R- and K-starting PANK3 variants, whereas the substrate-binding pocket mutants W247A, H255A, and D257A/D258A failed to rescue substrate degradation (Figures 8J and 8K). Likewise, wild-type RanBP10, but not the corresponding pocket mutants W135A, H143A, or D145A/D146A, restored degradation of these substrates (Figures S8G and S8H). Together, these biochemical, structural, and cellular findings establish the B30.2 domains of RanBP9 and RanBP10 as substrate-recognition modules that recognize both Met-X and canonical Arg/N-degrons. Thus, RanBP9 and RanBP10 define a previously unrecognized class of Arg/N-degron recognins that employ a substrate-recognition mechanism distinct from that of the canonical UBR family.

## Discussion

Cellular metabolism relies on precise control of rate-limiting enzymes, and PANK occupies a particularly important position in this regulatory network as the rate-limiting enzyme of CoA biosynthesis. Recent studies demonstrating that pharmacological activation of PANK1 by sodium-glucose cotransporter 2 inhibitors (SGLT2i) stimulates CoA synthesis, promotes metabolic fuel utilization, and enhances the contractility of human cardiomyocytes underscore the importance of PANK1 activity in coordinating cellular metabolic function.^54^ These findings further raise the fundamental question of whether PANK1 abundance, in addition to its enzymatic activity, is subject to regulated control. Whether ubiquitin-dependent proteolysis directly regulates PANK proteins to modulate CoA biosynthesis, however, has remained unknown. In this study, we identify PANK1β and PANK3 as previously unrecognized substrates of the mammalian Arg/N-degron pathway, establishing regulated protein degradation as an additional layer of CoA homeostasis. Mechanistically, we show that two distinct E3 ligase systems, the UBR4–KCMF1 heterodimeric ligase and the MKLN1–CTLH complex, independently recognize the same N-terminal MK Arg/N-degron to promote PANK degradation and thereby additively constrain cellular CoA synthesis. We further identify RanBP9 and RanBP10 as previously unrecognized classes of Arg/N-degron recognins that recognize both Met-X and canonical Arg/N-degrons through their B30.2 domains, revealing a mode of substrate recognition distinct from that of canonical UBR family proteins. Collectively, these findings expand the substrate-recognition landscape of the mammalian Arg/N-degron pathway and uncover regulated PANK degradation as a mechanism linking ubiquitin-dependent proteolysis to the control of CoA homeostasis. Cellular metabolism relies on precise control of rate-limiting enzymes, yet whether ubiquitin-dependent proteolysis directly regulates the first and rate-limiting step of CoA biosynthesis has remained unknown. In this study, we identify PANK1β and PANK3 as previously unrecognized substrates of the mammalian Arg/N-degron pathway, establishing regulated protein degradation as an additional layer of CoA homeostasis. Mechanistically, we show that two distinct E3 ligase systems, the UBR4-KCMF1 heterodimeric ligase and the MKLN1-CTLH complex, independently recognize the same N-terminal MK Arg/N-degron to promote PANK degradation and thereby additively constrain cellular CoA synthesis. We further identify RanBP9 and RanBP10 as a previously unrecognized class of Arg/N-degron recognins that recognize both Met-X and canonical Arg/N-degrons through their B30.2 domains, revealing a mode of substrate recognition distinct from that of canonical UBR family proteins. Collectively, these findings expand the substrate-recognition landscape of the mammalian Arg/N-degron pathway and uncover regulated PANK degradation as a mechanism linking ubiquitin-dependent proteolysis to the control of CoA homeostasis.

Although multiple mechanisms regulate CoA biosynthesis, including feedback inhibition by CoA thioesters, transcriptional regulation, and reversible phosphorylation,^2,9–13,18^ these mechanisms primarily modulate PANK activity or expression in response to metabolic demand. Our findings reveal an additional layer of regulation in which the cellular abundance of PANK proteins is actively controlled through selective ubiquitin-proteasome-dependent degradation. Whereas allosteric and post-translational mechanisms can rapidly tune the activity of pre-existing PANK proteins, regulated proteolysis controls the cellular abundance of these rate-limiting enzymes and thereby the capacity for CoA production. Consistent with this model, simultaneous disruption of the two E3 ligase systems markedly increased PANK abundance and expanded the intracellular pools of CoA and acetyl-CoA, demonstrating that PANK degradation has a substantial impact on CoA metabolism. Thus, ubiquitin-dependent proteolysis operates alongside metabolic, transcriptional, and post-translational mechanisms to maintain CoA homeostasis by controlling not only PANK activity but also the abundance of the enzymes that determine the entry point into CoA biosynthesis.

One of the most unexpected findings of this study is that two mechanistically distinct E3 ligase systems converge on the same N-terminal MK Arg/N-degron. In many ubiquitin-dependent degradation pathways, a given degron is recognized by a dedicated substrate receptor or by closely related paralogs.^31,55^ By contrast, UBR4–KCMF1 and the MKLN1-containing CTLH complex independently recognize the same degron while remaining genetically and biochemically independent. Re-expression of either ligase complex alone was sufficient to restore PANK degradation, whereas simultaneous disruption of both complexes resulted in substantially greater PANK stabilization than disruption of either complex alone, demonstrating that the two ligases act in parallel and contribute additively rather than through a hierarchical pathway.^56^ Such convergent degron recognition may provide robustness in the control of PANK turnover, ensuring that fluctuations in the activity of one E3 ligase do not completely abrogate substrate degradation. At the same time, the existence of two independent recognition systems raises the possibility that PANK degradation can be tuned according to distinct cellular contexts. Whether UBR4–KCMF1 and MKLN1-CTLH are differentially regulated by nutrient availability, metabolic stress, developmental programs, or tissue-specific signals remains an important question for future investigation.

Our findings substantially expand the substrate-recognition landscape of the mammalian Arg/N-degron pathway. Previous studies have established UBR family proteins, as well as the non-E3 autophagy regulator p62, as Arg/N-recognins that recognize destabilizing N-terminal residues, including canonical Arg/N-degrons and Met-Φ degrons.^31,45,46,48,57^ More recently, N-terminal methionine followed by lysine or arginine was proposed to constitute additional Arg/N-degrons recognized by UBR proteins.^47^ Here, we demonstrate that UBR4-KCMF1 complex recognizes the MK/MR Arg/N-degrons through the ZZ domain of KCMF1, establishing this domain as a substrate-recognition module for these previously expanded Arg/N-degrons. More importantly, we identify RanBP9 and RanBP10 as a distinct class of Arg/N-degron recognins within the CTLH complex. Unlike canonical Arg/N-recognins, which employ conserved UBR-box and ZZ domains for substrate recognition, RanBP9 and RanBP10 recognize both Met-X and canonical Arg/N-degrons through their B30.2 domains. Notably, the substrate specificity of RanBP9/10 only partially overlaps with that of UBR proteins, with RanBP9/10 recognizing a restricted subset of Met-X Arg/N-degrons while also engaging multiple canonical Arg/N-degrons. These observations indicate that mammalian Arg/N-degron recognition is mediated by multiple structurally distinct recognin systems with overlapping yet complementary substrate specificities, revealing a broader and more diverse architecture of the Arg/N-degron pathway than previously appreciated.

Our structural and biochemical analyses provide a molecular explanation for this alternative mode of degron recognition. The crystal structures, together with mutagenesis, peptide-binding, and cellular complementation experiments, demonstrate that the B30.2 domains of RanBP9 and RanBP10 directly engage N-terminal degrons through a compact surface pocket that is mechanistically distinct from the substrate-recognition interface characterized in UBR family proteins.^44,58,59^ Remarkably, the same B30.2 pocket accommodates both Met-X and canonical Arg/N-degrons, indicating an unexpected degree of structural plasticity in degron recognition. The shared use of conserved pocket residues by RanBP9 and RanBP10 further suggests that this recognition mechanism arose as a conserved feature of these paralogs. Given the broad distribution of B30.2 domains among metazoan proteins and their established roles in diverse protein-protein recognition events,^60^ it will be important to determine whether recognition of N-terminal degrons represents a more general function of B30.2-containing proteins beyond the CTLH complex.

Finally, our findings broaden both the substrate-recognition repertoire and the functional landscape of the CTLH E3 ligase complex. Previous studies have identified GID4, WDR26, and MKLN1 as substrate receptors involved in the Pro/N-degron pathway, metabolic regulation, and C-degron pathways, respectively.^20,32,36,39–43^ Our identification of RanBP9 and RanBP10 as Arg/N-degron recognins establishes the CTLH complex as a previously unrecognized branch of the mammalian Arg/N-degron pathway and reveals that a single E3 ligase assembly can engage distinct classes of degrons through multiple substrate-recognition modules. More broadly, the convergence of UBR4-KCMF1 and MKLN1-CTLH on PANK1β and PANK3 illustrates how parallel ubiquitin-dependent degradation pathways can provide robust control over the abundance of the rate-limiting metabolic enzymes. By linking the expanded Arg/N-degron recognition network to PANK turnover and CoA biosynthesis, our findings establish regulated proteolysis as an important component of CoA homeostasis and provide a framework for exploring how modulation of PANK stability or degron recognition may influence metabolic states in disease.

### Limitations of the study

This study establishes that UBR4–KCMF1 and the MKLN1-containing CTLH complex independently recognize a common MK Arg/N-degron to regulate PANK turnover and CoA biosynthesis. Nevertheless, several important questions remain. Although the two E3 ligase systems function additively, the physiological signals that regulate their activity and determine their relative contributions under different metabolic conditions remain unknown. In addition, while PANK1β and PANK3 are established here as physiological substrates of both ligases, the full repertoire of substrates recognized by RanBP9/10-mediated Arg/N-degron recognition remains to be defined. Finally, although our structural, biochemical, and cellular studies establish the molecular basis of degron recognition, determining how this newly identified branch of the Arg/N-degron pathway contributes to organismal physiology, metabolic adaptation, and disease will require future in vivo investigation.

## Methods

### Cell lines and cell culture

Human cell lines, including HEK293T and HeLa (ATCC) were maintained in Dulbecco’s modified Eagle’s Medium (Metacell) supplemented with 10% fetal bovine serum (FBS; Sigma-Aldrich) and 1% penicillin/streptomycin (Metacell). All cell line identities were authenticated by genotyping and routinely tested negative for mycoplasma contamination. Cells were cultured at 37 °C in a humidified incubator containing 5% CO2.

### Lentivirus production and transduction

Lentivirual particles were produced by co-transfecting HEK293T cells with packaging plasmids psPAX2 (Addgene), pMD2.G (Addgene), along with either LentiCRISPR v2 or pCDH expression constructs, using Liposomal Transfection Reagent (UElandy) according to the manufacturer’s instructions. Viral supernatants were harvested 48 h post-transfection, filtered through a 0.45 μm filter, and used to transduce recipient cells in the presence of 8 µg/ml polybrene. At 48 h post-transduction, cells were selected with puromycin (2µg/ml), blasticidin (10µg/ml) and/or hygromycin (200µg/ml) for 3-4 days to establish stable cell lines.

### Global protein stability (GPS) assay

Protein stability was measured using the Global Protein Stability (GPS) assay, a bicistronic reporter system in which green fluorescent protein (GFP) is fused to target proteins or peptide of interest, while red fluorescent protein (RFP) is expressed from the same transcript via an internal ribosome entry site (IRES), enabling normalization for transcriptional and translational variability. The GFP/RFP ratio serves as a quantitative readout of protein stability in living cells. Two reporter constructs were employed in this study. The pCDH-Ub-MCS-GFP-IRES-RFP vector expresses a Ub-target protein/fragment-GFP fusion protein, wherein cleavage of the ubiquitin moiety by endogenous deubiquitinating enzymes upon translation exposes the target protein or fragments at the N-terminus of GFP. The pHAGE-GFP-MCS-IRES-RFP vector, in contrast, generates a GFP-target protein/fragment fusion, placing the target at the C-terminus of GFP.

To establish stable reporter cell lines, lentiviral particles carrying the respective GPS reporters were produced and used to transduce HEK293T or HeLa cells. Transduced cells were selected with blasticidin (10μg/mL) to obtain stable populations. Subsequently, genetic manipulations, including overexpression or endogenous knockout of the gene of interest, were performed in these reporter lines, followed by selection with puromycin (2μg/mL) for 3 days to enrich for successfully modified cells.

Finally, protein or fragment stability was assessed by measuring the cellular GFP/RFP ratio via fluorescence-activated cell sorting (FACS), with RFP serving as an internal control. Data was collected by LSR Fortressa instrument (Becton Dickinson) and analyzed by FlowJo software v.10.

### CRISPR-Cas9 screen

CRISPR screens were performed using a custom gRNA library targeting protein degradation associated genes, comprising 11,180 guide RNAs (gRNAs) targeting 1,118 distinct human genes, along with 300 intergenic control gRNAs. Lentiviral particles were generated by co-transfecting HEK293T cells with the library plasmids, the packaging plasmid psPAX2, and the envelope plasmid pMD2.G using X-tremeGENE HP DNA transfection reagent (Sigma-Aldrich). Viral supernatants were harvested 48 h after transfection and used to transduce target cells at a multiplicity of infection (MOI) of 0.3 in the presence of 8μg/mL polybrene (Beyotime), while maintaining a coverage of 500 cells per gRNA. At 48h post-transduction, cells were selected with 2μg/mL puromycin until all untransduced cells were completely eliminated. Surviving cells were pooled and expanded for 10 days to allow CRISPR-Cas9-mediated gene knockout. All cells were harvested and subjected to FACS. The top 5% of cells exhibiting the highest GFP/RFP ratio were collected. To maintain adequate library representation, approximately 6×106 cells were sorted, corresponding to ∼500-fold gRNA coverage. An additional 6×106 cells from the remaining 95% of the population were retained as the control group.

Genomic DNA was extracted from the sorted and the control population using the TIANamp genomic DNA Kit (Tiangen). Integrated gRNA sequences were subsequently amplified from genomic DNA by PCR using Next Ultra II Q5 Master Mix (New England Biolabs). Each 50 µl PCR reaction contained 4µg genomic DNA, and sufficient parallel reactions were performed to recover genomic DNA representing at least 200-fold library coverage. PCR products from all reactions were pooled, purified and subjected to next-generation sequencing.

Sequencing reads were aligned to the reference gRNA library, and the gRNA abundance was quantified using MAGeCK to generate raw read counts for each sgRNA. A count matrix was constructed from the sorted and control populations, and differential gRNAs enrichment analysis was subsequently performed by MAGeCK to identify the genes whose targeting gRNAs were significantly enriched in the sorted population relative to the control population. MAGeCK output was visualized as a scatter plot, with genes arranged along the x axis and the corresponding −log10(MAGeCK positive selection score “pos|score”) plotted on the y axis.

### Generation of CRISPR–cas9 genome edited cell lines

For generation of clonal knockout cell lines, the gRNAs listed in the Key Resources Table were cloned into the pSpCas9 (BB)-2A-GFP (PX458) plasmid (Addgene). For generation of UBR4-ΔUBR mutant (Δ1653–1725), gRNAs were designed to introduce an in-frame deletion of amino acids 1653-1725. For transient transfection, target cells were seeded 18-24 h prior to transfection to achieve approximately 50-70% confluency at the time of transfection. PX458-gRNA plasmids were transiently transfected using Liposomal Transfection Reagent (UElandy) according to the manufacturer’s instructions. At 48 h post-transfection, cells were harvested and GFP positive cells were isolated by FACS at the single-cell level into 96-well plate to generate clonal knockout/mutant cell lines. Following expansion of the clones, total protein and genomic DNA were extracted from individual clones. Western blot analysis was performed to confirm loss of the target protein, and the genomic region spanning the gRNA target site was amplified by PCR and subjected to Sanger sequencing. Clones exhibiting complete loss of protein expression together with indel mutations at the target locus were selected for subsequent experiments.

### Immunoblotting

Cells were rinsed with PBS and lysed on ice for 20min in RIPA buffer (50mM Tris-HCl, pH7.4, 150mM NaCl, 2mM EDTA, 1% NP-40, 0.1% SDS) supplemented with 1mM PMSF and protease inhibitor cocktail (MCE). Lysates were sonicated (10% amplitude, 0.5s on/off pulses, 20 cycles), and protein concentrations were determined using Bradford Assay (Beyotime). Samples were denatured in 5x SDS-PAGE loading buffer (250mM Tris-HCl pH6.8, 10% SDS, 30% glycerol, 5% β-mercapitalethanol, 0.02% bromophenol blue) at 95°C for 5min. Equal amounts of protein (10-30 µg) were separated by SDS-PAGE and transferred onto a 0.45μm PVDF membrane (Millipore). Membranes were blocked with 5% non-fat milk in TBST for 1h at room temperature, then incubated with primary antibodies (1:1000) overnight at 4°C, followed by HRP-conjugated secondary antibody (1:10000) for 1h at room temperature. Protein bands were detected using Amersham ECL Detection Reagent (Cytiva) or Super ECL Detection Reagents (Yeasen) and visualized with Tanon 5200 Chemiluminescent Imaging System (Tanon).

### Co-Immunoprecipitation (Co-IP)

Cells were rinsed with PBS, harvested by scraping, and lysed in ice-cold lysis buffer (50mM Tris-HCl, pH 7.4, 250mM NaCl, 0.5% Triton X-100, 10% glycerol, 1mM DTT, 1 mM PMSF and protease inhibitors). An aliquot of the lysate was retained as the input control, and 1mg of the remaining lysate was incubated with GFP or HA-conjugated magnetic beads (Smart-Lifesciences) overnight at 4 °C with end-to-end rotation. The immunoprecipitates were washed three times with lysis buffer, and bound proteins were eluted in SDS loading buffer and denatured at 95 °C for 5min before immunoblot analysis.

### Protein expression and purification

For GST-tagged proteins, the gene fragments encoding wild-type or mutant regions of human KCMF1 (amino acids 1-70), RanBP9 (amino acids 108-350) and RanBP10 (amino acids 29-238) were cloned into the pGEX-6P-1 vector (Cytiva) via seamless cloning to generate N-terminal GST-fusion constructs. The resulting recombinant plasmids were transformed into E. coli Rosetta (DE3) cells (Sigma-Aldrich) and selected with ampicillin. Bacteria were cultured in Luria-Bertani (LB) medium at 37°C until OD600 reached 0.6-0.8, after which protein expression was induced with 0.2 mM isopropyl β-D-thiogalactopyranoside (IPTG) at 18°C for 16 h. Cells were harvested by centrifugation and resuspended in lysis buffer (50mM Tris-HCl pH 7.5, 150mM NaCl and 0.05% NP-40), followed by sonication on ice. Cell debris was removed by centrifugation at 12,000 rpm for 20min, and the clarified supernatant was then incubated with Glutathione Sepharose 4B resin at 4 °C for 4h. The resin was washed with 100 mM Tris-HCl (pH 8.0), and bound proteins were eluted by elution buffer (100mM Tris-HCl, pH 8.0 containing reduced glutathione). Purified proteins were stored at −80°C.

For His-SUMO fusion proteins, the human RanBP9 fragment (residues 145-350) was amplified with an N-terminal extension encoding either the MKLI or RKLI sequence and cloned into the pET28-MKH8SUMO vector (Addgene) by seamless cloning. The resulting construct encoded an N-terminal 8 x His-SUMO tag followed by a TEV (Tobacco Etch Virus) cleavage site (LYFQY↓M/RKLI), thereby exposing the M/RKLI sequence at the N-terminus after cleavage. The recombinant plasmids were expressed in E. coli Rosetta (DE3) cells with kanamycin selection. Cell culture, protein induction and cell harvesting were performed as described for GST fusion proteins. Cell pellets were resuspended in lysis buffer (20mM Tris-HCl pH 7.5, 400mM NaCl and 2mM β-mercaptoethanol) and lysed by sonication. The clarified lysate was incubated with Ni-NTA beads at 4°C for 1h. The beads were washed with washing buffer (20mM Tris-HCl pH 7.5, 400mM NaCl and 25mM imidazole pH 7.5), and bound proteins were eluted with elution buffer (20mM Tris-HCl pH 7.5, 400mM NaCl and 500mM imidazole pH 7.5). The eluates were dialyzed overnight at 4°C against dialysis buffer (20mM Tris-HCl pH 7.5, 300mM NaCl and 2mM β-mercaptoethanol) in the presence of TEV protease at a 1:30 molar ratio. The cleaved samples were subsequently reloaded onto a Ni-NTA column to remove the cleaved His-SUMO tag and TEV protease. The flow-through samples were further purified by anion-exchange chromatography on a HiTrap Q HP column (GE healthcare), followed by size-exclusion chromatography on a Superdex 200 Increase 10/300 GL (GE healthcare) equilibrated in gel-filtration buffer (20mM Tris-HCl pH 7.5, 150mM NaCl and 1mM DTT). Purified proteins were concentrated and stored at −80°C.

### Protein Crystallization

Crystallization trials were performed at 18°C using the sitting-drop vapor diffusion method. Briefly, 1μL of purified protein was mixed with 1μL of reservoir solution. Crystals of M/RKLI-fused RanBP9 (residues 145-350) were obtained in a reservoir solution containing 0.2M ammonium formate and 20% (w/v) polyethylene glycol 3,350. Before data collection, crystals were cryoprotected by soaking them in the corresponding reservoir solution supplemented with 20% (v/v) glycerol prior to flash-freezing in liquid nitrogen.

### Data collection and structure determination

Diffraction data were collected at beamline BL17U or BL19U of Shanghai Synchrotron Radiation Facility (SSRF) and processed with XDS. The structure of selenomethionine-substituted RanBP9 fragment (residues 108-350) was determined by the single-wavelength anomalous diffraction (SAD). Initial phases and model building were obtained using AutoSol and AutoBuild respectively, as implemented in PHENIX package. The resulting model was further refined by manual building in Coot. Structural figures were prepared with PyMOL (https://www.pymol.org/2/).

### Peptide pull-down assay

To analyze protein–peptide interactions,1µg of biotinylated peptides were incubated with 1-2µg of GST-fused proteins in binding buffer (50mM Tris-HCl 7.5, 250mM NaCl, 0.1% NP-40, 1mM PMSF) overnight at 4 °C. Streptavidin-conjugated magnetic beads (Smart-Lifesciences) were then added to the mixture, and incubated for an additional 1h with rotation to capture the biotinylated peptides and associated proteins. The beads were subsequently washed three times with binding buffer, and the bound proteins were eluted and analyzed by SDS-PAGE followed by Western blotting.

### UPLC-MS/MS

A total of 1×10^7^ cells were collected and resuspended in 400 μL of pre-cooled 80% methanol containing 1 μL HMG-CoA-d3 (1 mg/mL) as an internal standard. Cell suspensions were homogenized by ultrasonication (3 cycles of 3 s pulse) and then centrifuged to remove cellular debris. The resulting supernatant was mixed with 1 mL methyl tert-butyl ether (MTBE), vortexed for 10 min, and supplemented with 100 μL water, followed by an additional 2 min of vortexing. Samples were allowed to stand for 10 min at room temperature to facilitate phase separation and were subsequently centrifuged at 12,000 g for 10 min at 4 °C, resulting in the separation of the upper organic and lower aqueous phase. The aqueous phase was carefully collected, passed through a 0.22 μm centrifugal filter unit by centrifugation at 12,000×g for 5 min at 4 °C. Finally, 5 μL of the filtrate was injected into the ultra-performance liquid chromatography-tandem mass spectrometry (UPLC-MS/MS) system for metabolite analysis.

Chromatographic separation was performed on a BEH C18 column (1.7 μm, 2.1 mm × 100 mm) at a flow rate of 0.1 mL/min. The mobile phase consisted of water containing 10 mM ammonium acetate (solvent A, pH 9.0) and acetonitrile (ACN; solvent B). The gradient elution program was set as follows: 0−2 min, 2% B; 2−18 min, 2−14% B; 18−20 min, 14−95% B; 20−21 min, 95% B; 21−24 min, 95−2% B; and 24−25 min, 2% B.

Mass spectrometric analysis was performed on a 5500 QTRAPhybrid triple quadrupole-linear ion trap mass spectrometer (AB Sciex) equipped with a Turbo V ion spray electrospray ionization (ESI) source. Analytes were detected in multiple reaction monitoring (MRM) mode under both positive and negative ionization polarities, with a dwell time of 50 ms. Instrument parameters were optimized as follows: curtain gas (CUR), 40 psi; nebulizer gas (GS1), 30 psi; auxiliary gas (GS2), 30 psi; ion spray voltage (IS), 5,500 V; collision-activated dissociation gas (CAD), medium; and source temperature, 475 °C.

Raw UPLC-MS/MS data were processed with Analyst 1.7 (AB Sciex) and quantification was performed with MultiQuant 3.0.2 (AB Sciex). Metabolite concentrations were determined based on calibration curves generated using analytical standards with a purity of > 98%. The resulting concentrations were normalized to the aqueous-phase volume and cell number.

### Quantification and statistical analysis

No statistical methods were used to predetermine sample sizes, and no data were excluded from the analyses. Statistical details, including the number of biological replicates and statistical tests used, are provided in the corresponding figure legends. Quantitative data are presented as mean ± s.d. from at least two independent biological replicates. Statistical significance between two groups was assessed using an unpaired two-tailed Student’s t-test. For comparisons involving multiple groups, statistical significance was determined by analysis of variance (ANOVA) followed by Bonferroni post hoc correction for multiple comparisons. Statistical analyses were performed using Prism 10.1.2 (GraphPad). P values are reported in the figures and corresponding figure legends.

### Key resources table

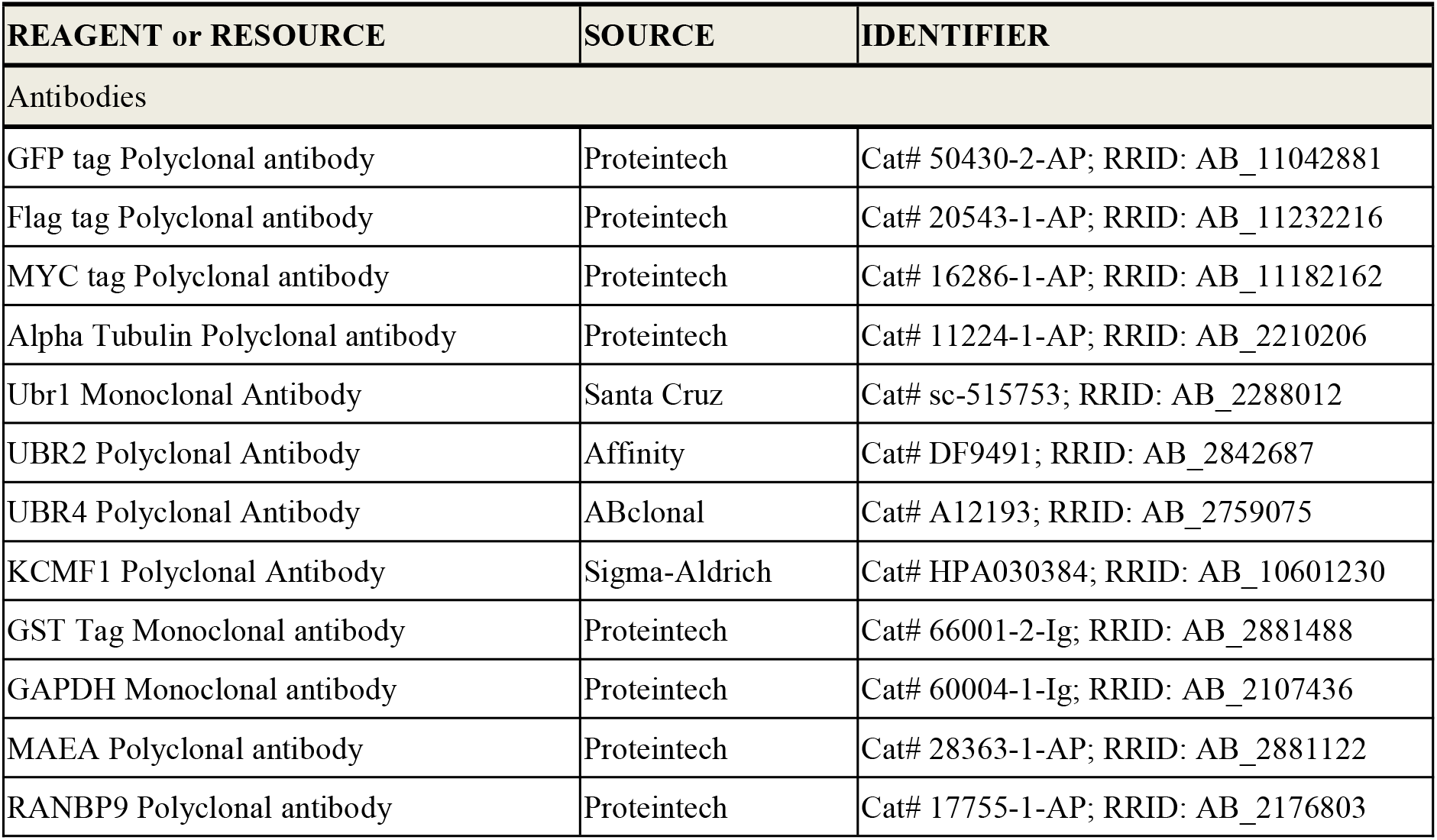
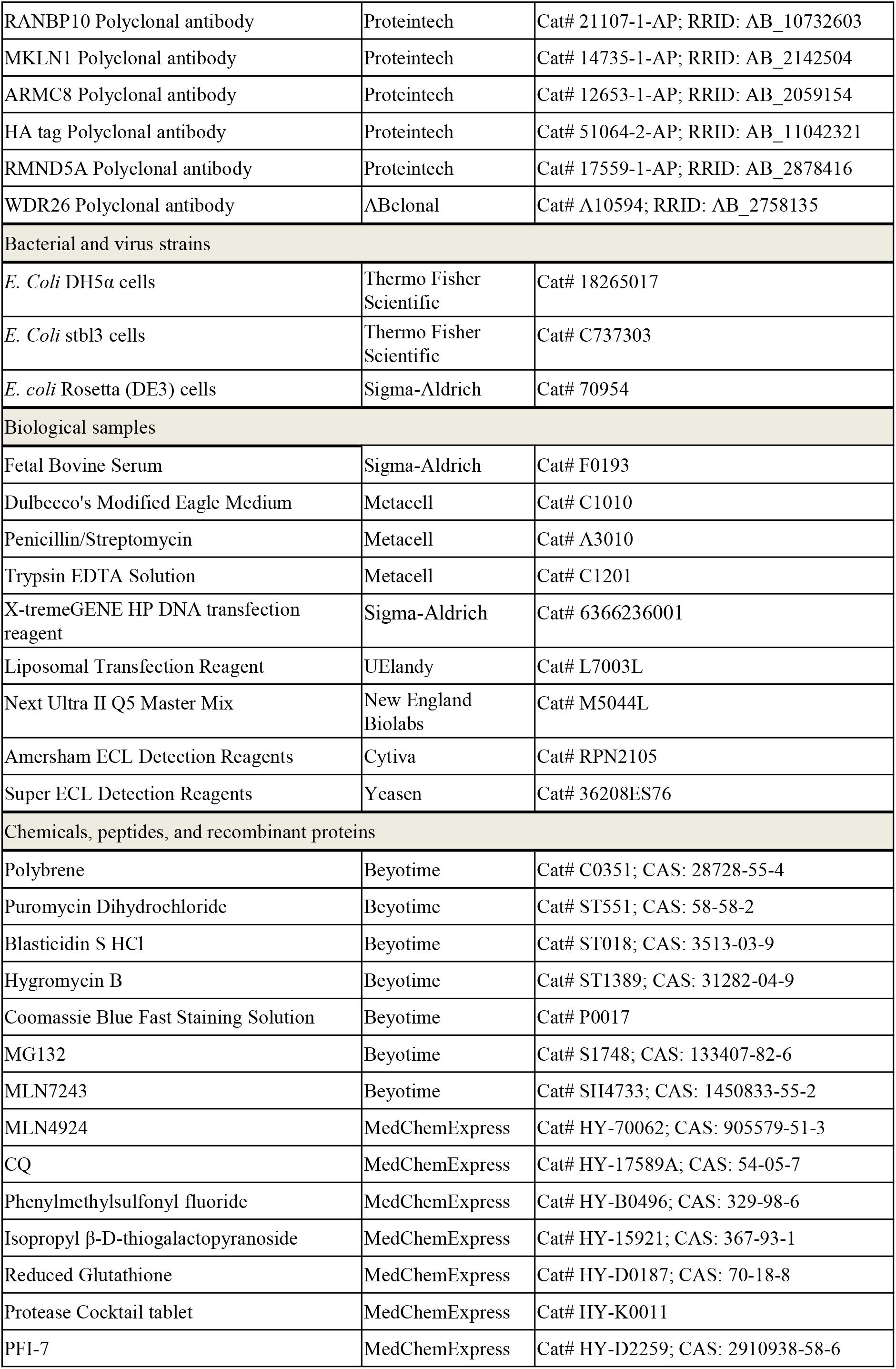
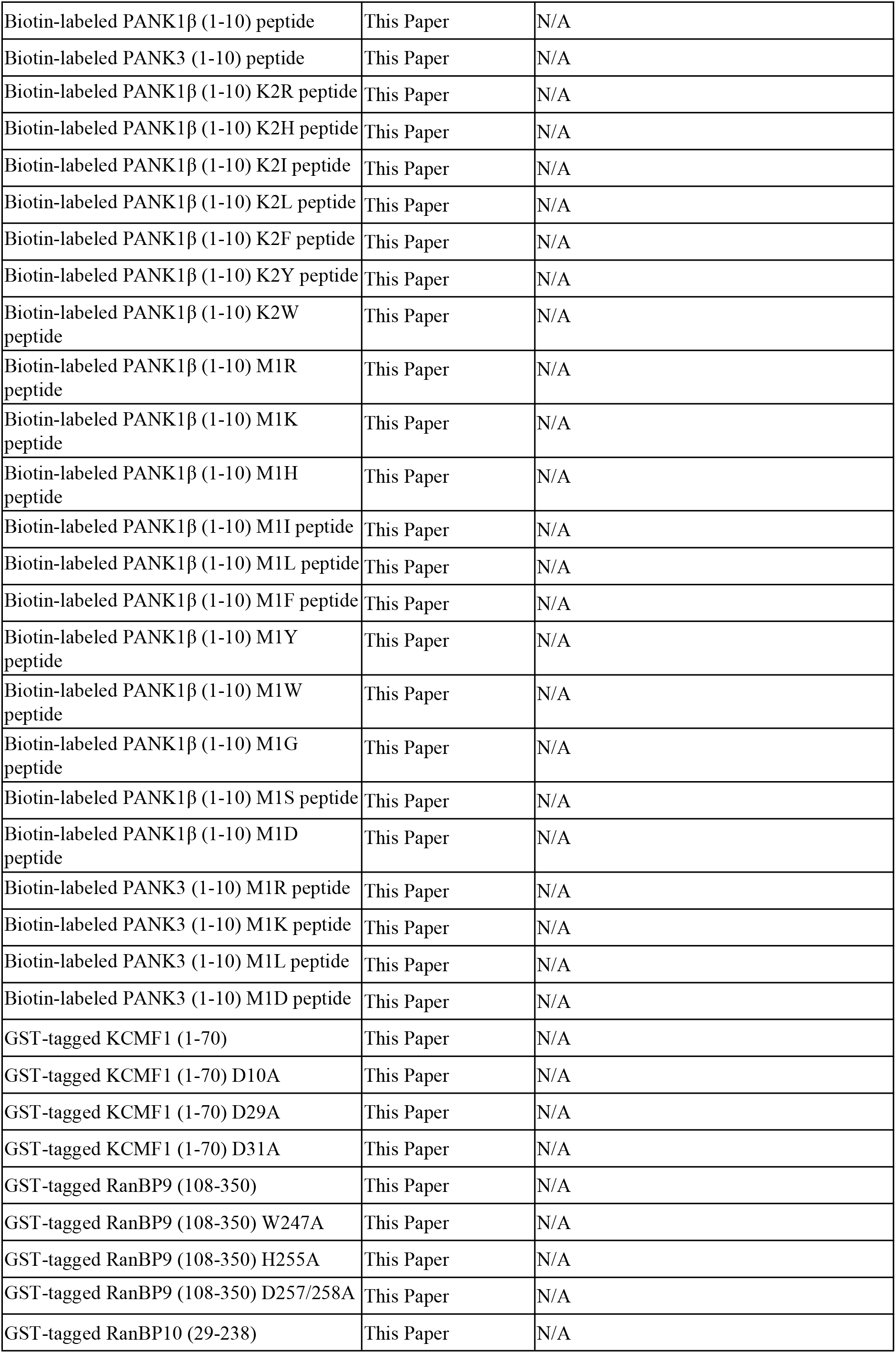
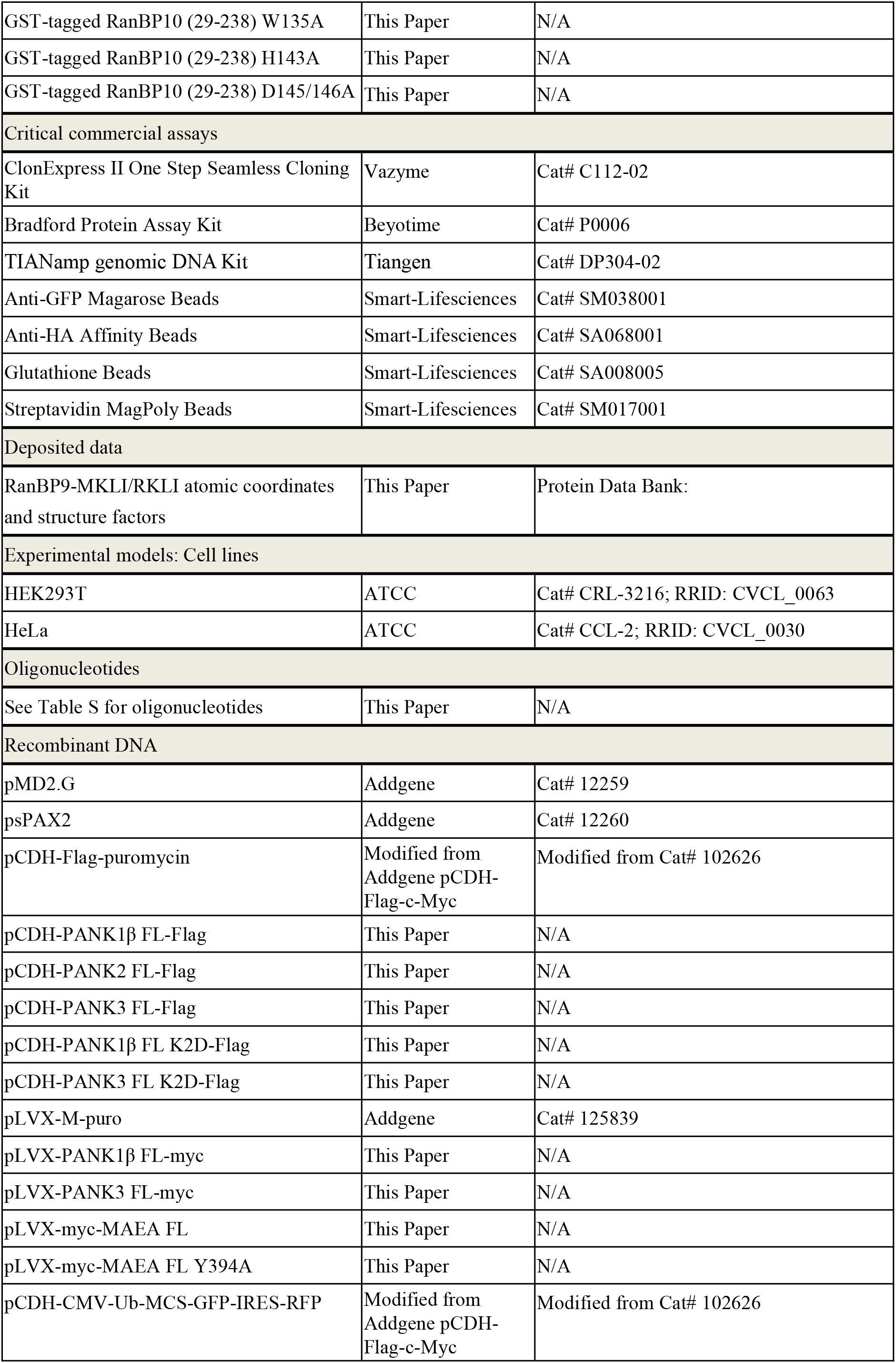
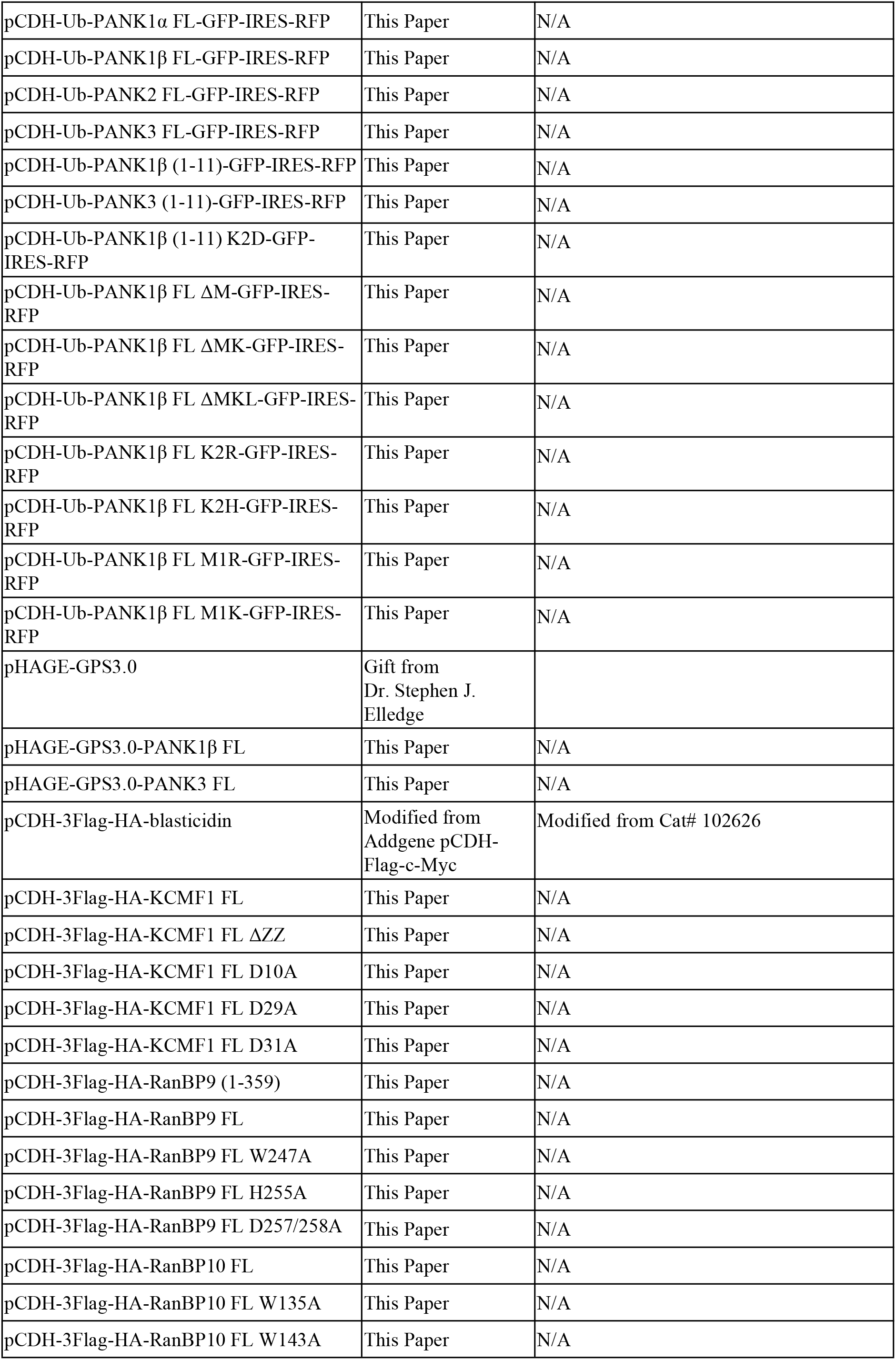
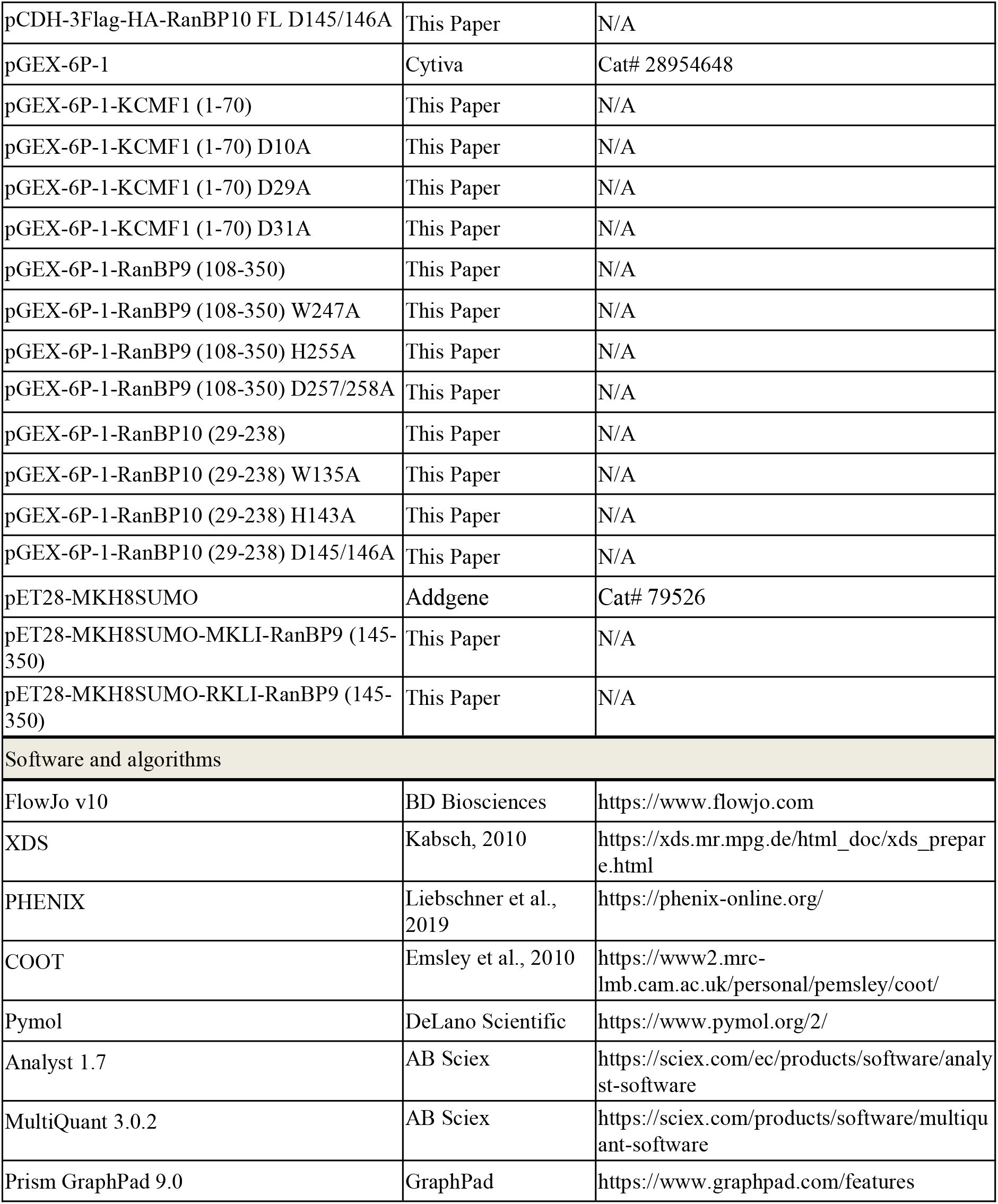

| REAGENT or RESOURCE | SOURCE | IDENTIFIER |
| --- | --- | --- |
| Antibodies |  |  |
| GFP tag Polyclonal antibody | Proteintech | Cat# 50430-2-AP; RRID: AB_11042881 |
| Flag tag Polyclonal antibody | Proteintech | Cat# 20543-1-AP; RRID: AB_11232216 |
| MYC tag Polyclonal antibody | Proteintech | Cat# 16286-1-AP; RRID: AB_11182162 |
| Alpha Tubulin Polyclonal antibody | Proteintech | Cat# 11224-1-AP; RRID: AB_2210206 |
| Ubr1 Monoclonal Antibody | Santa Cruz | Cat# sc-515753; RRID: AB_2288012 |
| UBR2 Polyclonal Antibody | Affinity | Cat# DF9491; RRID: AB_2842687 |
| UBR4 Polyclonal Antibody | ABclonal | Cat# A12193; RRID: AB_2759075 |
| KCMF1 Polyclonal Antibody | Sigma-Aldrich | Cat# HPA030384; RRID: AB_10601230 |
| GST Tag Monoclonal antibody | Proteintech | Cat# 66001-2-Ig; RRID: AB_2881488 |
| GAPDH Monoclonal antibody | Proteintech | Cat# 60004-1-Ig; RRID: AB_2107436 |
| MAEA Polyclonal antibody | Proteintech | Cat# 28363-1-AP; RRID: AB_2881122 |
| RANBP9 Polyclonal antibody | Proteintech | Cat# 17755-1-AP; RRID: AB_2176803 |
| RANBP10 Polyclonal antibody | Proteintech | Cat# 21107-1-AP; RRID: AB_10732603 |
| MKLN1 Polyclonal antibody | Proteintech | Cat# 14735-1-AP; RRID: AB_2142504 |
| ARMC8 Polyclonal antibody | Proteintech | Cat# 12653-1-AP; RRID: AB_2059154 |
| HA tag Polyclonal antibody | Proteintech | Cat# 51064-2-AP; RRID: AB_11042321 |
| RMND5A Polyclonal antibody | Proteintech | Cat# 17559-1-AP; RRID: AB_2878416 |
| WDR26 Polyclonal antibody | ABclonal | Cat# A10594; RRID: AB_2758135 |
| Bacterial and virus strains |  |  |
| *E. Coli* DH5α cells | Thermo Fisher Scientific | Cat# 18265017 |
| *E. Coli* stbl3 cells | Thermo Fisher Scientific | Cat# C737303 |
| *E. coli* Rosetta (DE3) cells | Sigma-Aldrich | Cat# 70954 |
| Biological samples |  |  |
| Fetal Bovine Serum | Sigma-Aldrich | Cat# F0193 |
| Dulbecco's Modified Eagle Medium | Metacell | Cat# C1010 |
| Penicillin/Streptomycin | Metacell | Cat# A3010 |
| Trypsin EDTA Solution | Metacell | Cat# C1201 |
| X-tremeGENE HP DNA transfection reagent | Sigma-Aldrich | Cat# 6366236001 |
| Liposomal Transfection Reagent | Uelandy | Cat# L7003L |
| Next Ultra II Q5 Master Mix | New England Biolabs | Cat# M5044L |
| Amersham ECL Detection Reagents | Cytiva | Cat# RPN2105 |
| Super ECL Detection Reagents | Yeaden | Cat# 36208ES76 |
| Chemicals, peptides, and recombinant proteins |  |  |
| Polybrene | Beyotime | Cat# C0351; CAS: 28728-55-4 |
| Puromycin Dihydrochloride | Beyotime | Cat# ST551; CAS: 58-58-2 |
| Blasticidin S HCl | Beyotime | Cat# ST018; CAS: 3513-03-9 |
| Hygromycin B | Beyotime | Cat# ST1389; CAS: 31282-04-9 |
| Coomassie Blue Fast Staining Solution | Beyotime | Cat# P0017 |
| MG132 | Beyotime | Cat# S1748; CAS: 133407-82-6 |
| MLN7243 | Beyotime | Cat# SH4733; CAS: 1450833-55-2 |
| MLN4924 | MedChemExpress | Cat# HY-70062; CAS: 905579-51-3 |
| CQ | MedChemExpress | Cat# HY-17589A; CAS: 54-05-7 |
| Phenylmethylsulfonyl fluoride | MedChemExpress | Cat# HY-B0496; CAS: 329-98-6 |
| Isopropyl β-D-thiogalactopyranoside | MedChemExpress | Cat# HY-15921; CAS: 367-93-1 |
| Reduced Glutathione | MedChemExpress | Cat# HY-D0187; CAS: 70-18-8 |
| Protease Cocktail tablet | MedChemExpress | Cat# HY-K0011 |
| PFI-7 | MedChemExpress | Cat# HY-D2259; CAS: 2910938-58-6 |
| Biotin-labeled PANK1β (1-10) peptide | This Paper | N/A |
| Biotin-labeled PANK3 (1-10) peptide | This Paper | N/A |
| Biotin-labeled PANK1β (1-10) K2R peptide | This Paper | N/A |
| Biotin-labeled PANK1β (1-10) K2H peptide | This Paper | N/A |
| Biotin-labeled PANK1β (1-10) K2I peptide | This Paper | N/A |
| Biotin-labeled PANK1β (1-10) K2L peptide | This Paper | N/A |
| Biotin-labeled PANK1β (1-10) K2F peptide | This Paper | N/A |
| Biotin-labeled PANK1β (1-10) K2Y peptide | This Paper | N/A |
| Biotin-labeled PANK1β (1-10) K2W peptide | This Paper | N/A |
| Biotin-labeled PANK1β (1-10) M1R peptide | This Paper | N/A |
| Biotin-labeled PANK1β (1-10) M1K peptide | This Paper | N/A |
| Biotin-labeled PANK1β (1-10) M1H peptide | This Paper | N/A |
| Biotin-labeled PANK1β (1-10) M1I peptide | This Paper | N/A |
| Biotin-labeled PANK1β (1-10) M1L peptide | This Paper | N/A |
| Biotin-labeled PANK1β (1-10) M1F peptide | This Paper | N/A |
| Biotin-labeled PANK1β (1-10) M1Y peptide | This Paper | N/A |
| Biotin-labeled PANK1β (1-10) M1W peptide | This Paper | N/A |
| Biotin-labeled PANK1β (1-10) M1G peptide | This Paper | N/A |
| Biotin-labeled PANK1β (1-10) M1S peptide | This Paper | N/A |
| Biotin-labeled PANK1β (1-10) M1D peptide | This Paper | N/A |
| Biotin-labeled PANK3 (1-10) M1R peptide | This Paper | N/A |
| Biotin-labeled PANK3 (1-10) M1K peptide | This Paper | N/A |
| Biotin-labeled PANK3 (1-10) M1L peptide | This Paper | N/A |
| Biotin-labeled PANK3 (1-10) M1D peptide | This Paper | N/A |
| GST-tagged KCMF1 (1-70) | This Paper | N/A |
| GST-tagged KCMF1 (1-70) D10A | This Paper | N/A |
| GST-tagged KCMF1 (1-70) D29A | This Paper | N/A |
| GST-tagged KCMF1 (1-70) D31A | This Paper | N/A |
| GST-tagged RanBP9 (108-350) | This Paper | N/A |
| GST-tagged RanBP9 (108-350) W247A | This Paper | N/A |
| GST-tagged RanBP9 (108-350) H255A | This Paper | N/A |
| GST-tagged RanBP9 (108-350) D257/258A | This Paper | N/A |
| GST-tagged RanBP10 (29-238) | This Paper | N/A |
| GST-tagged RanBP10 (29-238) W135A | This Paper | N/A |
| GST-tagged RanBP10 (29-238) H143A | This Paper | N/A |
| GST-tagged RanBP10 (29-238) D145/146A | This Paper | N/A |
| Critical commercial assays |  |  |
| ClonExpress II One Step Seamless Cloning Kit | Vazyme | Cat# C112-02 |
| Bradford Protein Assay Kit | Beyotime | Cat# P0006 |
| TIANamp genomic DNA Kit | Tiagen | Cat# DP304-02 |
| Anti-GFP Magarose Beads | Smart-Lifesciences | Cat# SM038001 |
| Anti-HA Affinity Beads | Smart-Lifesciences | Cat# SA068001 |
| Glutathione Beads | Smart-Lifesciences | Cat# SA008005 |
| Streptavidin MagPoly Beads | Smart-Lifesciences | Cat# SM017001 |
| Deposited data |  |  |
| RanBP9-MKLI/RKLI atomic coordinates and structure factors | This Paper | Protein Data Bank: |
| Experimental models: Cell lines |  |  |
| HEK293T | ATCC | Cat# CRL-3216; RRID: CVCL_0063 |
| HeLa | ATCC | Cat# CCL-2; RRID: CVCL_0030 |
| Oligonucleotides |  |  |
| See Table S for oligonucleotides | This Paper | N/A |
| Recombinant DNA |  |  |
| pMD2.G | Addgene | Cat# 12259 |
| psPAX2 | Addgene | Cat# 12260 |
| pCDH-Flag-puromycin | Modified from Addgene pCDH-Flag-c-Myc | Modified from Cat# 102626 |
| pCDH-PANK1β FL-Flag | This Paper | N/A |
| pCDH-PANK2 FL-Flag | This Paper | N/A |
| pCDH-PANK3 FL-Flag | This Paper | N/A |
| pCDH-PANK1β FL K2D-Flag | This Paper | N/A |
| pCDH-PANK3 FL K2D-Flag | This Paper | N/A |
| pLVX-M-puro | Addgene | Cat# 125839 |
| pLVX-PANK1β FL-myc | This Paper | N/A |
| pLVX-PANK3 FL-myc | This Paper | N/A |
| pLVX-myc-MAEA FL | This Paper | N/A |
| pLVX-myc-MAEA FL Y394A | This Paper | N/A |
| pCDH-CMV-Ub-MCS-GFP-IRES-RFP | Modified from Addgene pCDH-Flag-c-Myc | Modified from Cat# 102626 |
| pCDH-Ub-PANK1α FL-GFP-IRES-RFP | This Paper | N/A |
| pCDH-Ub-PANK1β FL-GFP-IRES-RFP | This Paper | N/A |
| pCDH-Ub-PANK2 FL-GFP-IRES-RFP | This Paper | N/A |
| pCDH-Ub-PANK3 FL-GFP-IRES-RFP | This Paper | N/A |
| pCDH-Ub-PANK1β (1-11)-GFP-IRES-RFP | This Paper | N/A |
| pCDH-Ub-PANK3 (1-11)-GFP-IRES-RFP | This Paper | N/A |
| pCDH-Ub-PANK1β (1-11) K2D-GFP-IRES-RFP | This Paper | N/A |
| pCDH-Ub-PANK1β FL ΔM-GFP-IRES-RFP | This Paper | N/A |
| pCDH-Ub-PANK1β FL ΔMK-GFP-IRES-RFP | This Paper | N/A |
| pCDH-Ub-PANK1β FL ΔMKL-GFP-IRES-RFP | This Paper | N/A |
| pCDH-Ub-PANK1β FL K2R-GFP-IRES-RFP | This Paper | N/A |
| pCDH-Ub-PANK1β FL K2H-GFP-IRES-RFP | This Paper | N/A |
| pCDH-Ub-PANK1β FL M1R-GFP-IRES-RFP | This Paper | N/A |
| pCDH-Ub-PANK1β FL M1K-GFP-IRES-RFP | This Paper | N/A |
| pHAGE-GPS3.0 | Gift from Dr. Stephen J. Elledge |  |
| pHAGE-GPS3.0-PANK1β FL | This Paper | N/A |
| pHAGE-GPS3.0-PANK3 FL | This Paper | N/A |
| pCDH-3Flag-HA-blasticidin | Modified from Addgene pCDH-Flag-c-Myc | Modified from Cat# 102626 |
| pCDH-3Flag-HA-KCMF1 FL | This Paper | N/A |
| pCDH-3Flag-HA-KCMF1 FL ΔZZ | This Paper | N/A |
| pCDH-3Flag-HA-KCMF1 FL D10A | This Paper | N/A |
| pCDH-3Flag-HA-KCMF1 FL D29A | This Paper | N/A |
| pCDH-3Flag-HA-KCMF1 FL D31A | This Paper | N/A |
| pCDH-3Flag-HA-RanBP9 (1-359) | This Paper | N/A |
| pCDH-3Flag-HA-RanBP9 FL | This Paper | N/A |
| pCDH-3Flag-HA-RanBP9 FL W247A | This Paper | N/A |
| pCDH-3Flag-HA-RanBP9 FL H255A | This Paper | N/A |
| pCDH-3Flag-HA-RanBP9 FL D257/258A | This Paper | N/A |
| pCDH-3Flag-HA-RanBP10 FL | This Paper | N/A |
| pCDH-3Flag-HA-RanBP10 FL W135A | This Paper | N/A |
| pCDH-3Flag-HA-RanBP10 FL W143A | This Paper | N/A |
| pCDH-3Flag-HA-RanBP10 FL D145/146A | This Paper | N/A |
| pGEX-6P-1 | Cytiva | Cat# 28954648 |
| pGEX-6P-1-KCMF1 (1-70) | This Paper | N/A |
| pGEX-6P-1-KCMF1 (1-70) D10A | This Paper | N/A |
| pGEX-6P-1-KCMF1 (1-70) D29A | This Paper | N/A |
| pGEX-6P-1-KCMF1 (1-70) D31A | This Paper | N/A |
| pGEX-6P-1-RanBP9 (108-350) | This Paper | N/A |
| pGEX-6P-1-RanBP9 (108-350) W247A | This Paper | N/A |
| pGEX-6P-1-RanBP9 (108-350) H255A | This Paper | N/A |
| pGEX-6P-1-RanBP9 (108-350) D257/258A | This Paper | N/A |
| pGEX-6P-1-RanBP10 (29-238) | This Paper | N/A |
| pGEX-6P-1-RanBP10 (29-238) W135A | This Paper | N/A |
| pGEX-6P-1-RanBP10 (29-238) H143A | This Paper | N/A |
| pGEX-6P-1-RanBP10 (29-238) D145/146A | This Paper | N/A |
| pET28-MKH8SUMO | Addgene | Cat# 79526 |
| pET28-MKH8SUMO-MKLI-RanBP9 (145-350) | This Paper | N/A |
| pET28-MKH8SUMO-RKLI-RanBP9 (145-350) | This Paper | N/A |
| Software and algorithms |  |  |
| FlowJo v10 | BD Biosciences | https://www.flowjo.com |
| XDS | Kabsch, 2010 | https://xds.mr.mpg.de/html_doc/xds_prepare.html |
| PHENIX | Liebschner et al., 2019 | https://phenix-online.org/ |
| COOT | Emsley et al., 2010 | https://www2.mrc-lmb.cam.ac.uk/personal/pemsley/coot/ |
| Pymol | DeLano Scientific | https://www.pymol.org/2/ |
| Analyst 1.7 | AB Sciex | https://sciex.com/ec/products/software/analyst-software |
| MultiQuant 3.0.2 | AB Sciex | https://sciex.com/products/software/multiquant-software |
| Prism GraphPad 9.0 | GraphPad | https://www.graphpad.com/features |

## Acknowledgements

We thank members of the Mi lab for helpful suggestions. This work was supported by the National Natural Science Foundation of China grants (82372678 and 82674261 to W.Y.M.), Research Foundation of Tianjin Municipal Education Commission grant (2025ZD005 to W.Y.M.), Key Laboratory of Breast Cancer Prevention and Therapy (Tianjin Medical University, Ministry of Education), State Key Laboratory of Druggability Evaluation and Systematic Translational Medicine (Tianjin Medical University Cancer Institute & Hospital, No. QZKF24-4), and Core Facility of Research Center of Basic Medical Sciences at Tianjin Medical University.

## Author contributions

W.Y.M. and Y.Y. designed and conceived the project. X.L.W., B.J.W., and L.Z. performed most of biochemical, structural, cellular, and metabolic experiments and prepared the figures. S.X.Y. and N.X.Z. carried out the GPS construction and assays. Y.L.Z. and F.Y.S. performed mechanistic studies of PANK degradation by UBR4-KCMF1. S.Q.Y. conducted molecular cloning and protein purification. W.Y.M., X.L.W., and C.D. wrote the manuscript with critical input from all authors. Y.Y., D.X.C., and C.D. reviewed and edited the manuscript. All authors read and approved the final manuscript.

## References

1. Sibon, O.C., and Strauss, E. (2016). Coenzyme A: to make it or uptake it? Nat Rev Mol Cell Biol 17, 605–606. 10.1038/nrm.2016.110.

2. Barritt, S.A., DuBois-Coyne, S.E., and Dibble, C.C. (2024). Coenzyme A biosynthesis: mechanisms of regulation, function and disease. Nat Metab 6, 1008–1023. 10.1038/s42255-024-01059-y.

3. Robishaw, J.D., and Neely, J.R. (1985). Coenzyme A metabolism. Am J Physiol 248, E1–9. 10.1152/ajpendo.1985.248.1.E1.

4. Rock, C.O., Calder, R.B., Karim, M.A., and Jackowski, S. (2000). Pantothenate kinase regulation of the intracellular concentration of coenzyme A. J Biol Chem 275, 1377–1383. 10.1074/jbc.275.2.1377.

5. Zhou, B., Westaway, S.K., Levinson, B., Johnson, M.A., Gitschier, J., and Hayflick, S.J. (2001). A novel pantothenate kinase gene (PANK2) is defective in Hallervorden-Spatz syndrome. Nat Genet 28, 345–349. 10.1038/ng572.

6. Rock, C.O., Karim, M.A., Zhang, Y.M., and Jackowski, S. (2002). The murine pantothenate kinase (Pank1) gene encodes two differentially regulated pantothenate kinase isozymes. Gene 291, 35–43. 10.1016/s0378-1119(02)00564-4.

7. Martinez, D.L., Tsuchiya, Y., and Gout, I. (2014). Coenzyme A biosynthetic machinery in mammalian cells. Biochem Soc Trans 42, 1112–1117. 10.1042/BST20140124.

8. Alfonso-Pecchio, A., Garcia, M., Leonardi, R., and Jackowski, S. (2012). Compartmentalization of mammalian pantothenate kinases. PLoS One 7, e49509. 10.1371/journal.pone.0049509.

9. Abiko, Y., Ashida, S.I., and Shimizu, M. (1972). Purification and properties of D-pantothenate kinase from rat liver. Biochim Biophys Acta 268, 364–372. 10.1016/0005-2744(72)90331-2.

10. Karasawa, T., Yoshida, K., Furukawa, K., and Hosoki, K. (1972). Feedback inhibition of pantothenate kinase by coenzyme A and possible role of the enzyme for the regulation of cellular coenzyme A level. J Biochem 71, 1065–1067. 10.1093/oxfordjournals.jbchem.a129854.

11. Zhang, Y.M., Rock, C.O., and Jackowski, S. (2005). Feedback regulation of murine pantothenate kinase 3 by coenzyme A and coenzyme A thioesters. J Biol Chem 280, 32594–32601. 10.1074/jbc.M506275200.

12. Vallari, D.S., Jackowski, S., and Rock, C.O. (1987). Regulation of pantothenate kinase by coenzyme A and its thioesters. J Biol Chem 262, 2468–2471.

13. Fisher, M.N., Robishaw, J.D., and Neely, J.R. (1985). The properties and regulation of pantothenate kinase from rat heart. J Biol Chem 260, 15745–15751.

14. Jiang, L., Kon, N., Li, T., Wang, S.J., Su, T., Hibshoosh, H., Baer, R., and Gu, W. (2015). Ferroptosis as a p53-mediated activity during tumour suppression. Nature 520, 57–62. 10.1038/nature14344.

15. Wang, S.J., Yu, G., Jiang, L., Li, T., Lin, Q., Tang, Y., and Gu, W. (2013). p53-Dependent regulation of metabolic function through transcriptional activation of pantothenate kinase-1 gene. Cell Cycle 12, 753–761. 10.4161/cc.23597.

16. Yang, L., Zhang, B., Wang, X., Liu, Z., Li, J., Zhang, S., Gu, X., Jia, M., Guo, H., Feng, N., et al. (2020). P53/PANK1/miR-107 signalling pathway spans the gap between metabolic reprogramming and insulin resistance induced by high-fat diet. J Cell Mol Med 24, 3611–3624. 10.1111/jcmm.15053.

17. Bohlig, L., Friedrich, M., and Engeland, K. (2011). p53 activates the PANK1/miRNA-107 gene leading to downregulation of CDK6 and p130 cell cycle proteins. Nucleic Acids Res 39, 440–453. 10.1093/nar/gkq796.

18. Dibble, C.C., Barritt, S.A., Perry, G.E., Lien, E.C., Geck, R.C., DuBois-Coyne, S.E., Bartee, D., Zengeya, T.T., Cohen, E.B., Yuan, M., et al. (2022). PI3K drives the de novo synthesis of coenzyme A from vitamin B5. Nature 608, 192–198. 10.1038/s41586-022-04984-8.

19. Chen, S.J., Wu, X., Wadas, B., Oh, J.H., and Varshavsky, A. (2017). An N-end rule pathway that recognizes proline and destroys gluconeogenic enzymes. Science 355. 10.1126/science.aal3655.

20. Yi, S.A., Sepic, S., Schulman, B.A., Ordureau, A., and An, H. (2024). mTORC1-CTLH E3 ligase regulates the degradation of HMG-CoA synthase 1 through the Pro/N-degron pathway. Mol Cell 84, 2166–2184 e2169. 10.1016/j.molcel.2024.04.026.

21. Hershko, A., Ciechanover, A., and Varshavsky, A. (2000). Basic Medical Research Award. The ubiquitin system. Nature medicine 6, 1073–1081. 10.1038/80384.

22. Dikic, I., and Schulman, B.A. (2023). An expanded lexicon for the ubiquitin code. Nature reviews. Molecular cell biology 24, 273–287. 10.1038/s41580-022-00543-1.

23. Varshavsky, A. (2019). N-degron and C-degron pathways of protein degradation. Proceedings of the National Academy of Sciences of the United States of America 116, 358–366. 10.1073/pnas.1816596116.

24. Varshavsky, A. (2014). Discovery of the biology of the ubiquitin system. Jama 311, 1969–1970. 10.1001/jama.2014.5549.

25. Ohsumi, Y. (2014). Historical landmarks of autophagy research. Cell research 24, 9–23. 10.1038/cr.2013.169.

26. Zheng, N., and Shabek, N. (2017). Ubiquitin Ligases: Structure, Function, and Regulation. Annual review of biochemistry 86, 129–157. 10.1146/annurev-biochem-060815-014922.

27. Sherpa, D., Chrustowicz, J., and Schulman, B.A. (2022). How the ends signal the end: Regulation by E3 ubiquitin ligases recognizing protein termini. Molecular cell 82, 1424–1438. 10.1016/j.molcel.2022.02.004.

28. Varshavsky, A. (1991). Naming a targeting signal. Cell 64, 13–15. 10.1016/0092-8674(91)90202-a.

29. Bachmair, A., and Varshavsky, A. (1989). The degradation signal in a short-lived protein. Cell 56, 1019–1032. 10.1016/0092-8674(89)90635-1.

30. Bachmair, A., Finley, D., and Varshavsky, A. (1986). In vivo half-life of a protein is a function of its amino-terminal residue. Science 234, 179–186. 10.1126/science.3018930.

31. Varshavsky, A. (2024). N-degron pathways. Proc Natl Acad Sci U S A 121, e2408697121. 10.1073/pnas.2408697121.

32. Hammerle, M., Bauer, J., Rose, M., Szallies, A., Thumm, M., Dusterhus, S., Mecke, D., Entian, K.D., and Wolf, D.H. (1998). Proteins of newly isolated mutants and the amino-terminal proline are essential for ubiquitin-proteasome-catalyzed catabolite degradation of fructose-1,6-bisphosphatase of Saccharomyces cerevisiae. J Biol Chem 273, 25000–25005. 10.1074/jbc.273.39.25000.

33. Maitland, M.E.R., Lajoie, G.A., Shaw, G.S., and Schild-Poulter, C. (2022). Structural and Functional Insights into GID/CTLH E3 Ligase Complexes. Int J Mol Sci 23. 10.3390/ijms23115863.

34. Liu, H., and Pfirrmann, T. (2019). The Gid-complex: an emerging player in the ubiquitin ligase league. Biol Chem 400, 1429–1441. 10.1515/hsz-2019-0139.

35. Qiao, S., Langlois, C.R., Chrustowicz, J., Sherpa, D., Karayel, O., Hansen, F.M., Beier, V., von Gronau, S., Bollschweiler, D., Schafer, T., et al. (2020). Interconversion between Anticipatory and Active GID E3 Ubiquitin Ligase Conformations via Metabolically Driven Substrate Receptor Assembly. Mol Cell 77, 150–163 e159. 10.1016/j.molcel.2019.10.009.

36. Mohamed, W.I., Park, S.L., Rabl, J., Leitner, A., Boehringer, D., and Peter, M. (2021). The human GID complex engages two independent modules for substrate recruitment. EMBO Rep 22, e52981. 10.15252/embr.202152981.

37. Sherpa, D., Chrustowicz, J., Qiao, S., Langlois, C.R., Hehl, L.A., Gottemukkala, K.V., Hansen, F.M., Karayel, O., von Gronau, S., Prabu, J.R., et al. (2021). GID E3 ligase supramolecular chelate assembly configures multipronged ubiquitin targeting of an oligomeric metabolic enzyme. Mol Cell 81, 2445–2459 e2413. 10.1016/j.molcel.2021.03.025.

38. van Gen Hassend, P.M., Pottikkadavath, A., Delto, C., Kuhn, M., Endres, M., Schonemann, L., and Schindelin, H. (2023). RanBP9 controls the oligomeric state of CTLH complex assemblies. J Biol Chem 299, 102869. 10.1016/j.jbc.2023.102869.

39. Chen, S.J., Kim, L., Song, H.K., and Varshavsky, A. (2021). Aminopeptidases trim Xaa-Pro proteins, initiating their degradation by the Pro/N-degron pathway. Proc Natl Acad Sci U S A 118. 10.1073/pnas.2115430118.

40. Lampert, F., Stafa, D., Goga, A., Soste, M.V., Gilberto, S., Olieric, N., Picotti, P., Stoffel, M., and Peter, M. (2018). The multi-subunit GID/CTLH E3 ubiquitin ligase promotes cell proliferation and targets the transcription factor Hbp1 for degradation. Elife 7. 10.7554/eLife.35528.

41. Gottemukkala, K.V., Chrustowicz, J., Sherpa, D., Sepic, S., Vu, D.T., Karayel, O., Papadopoulou, E.C., Gross, A., Schorpp, K., von Gronau, S., et al. (2024). Non-canonical substrate recognition by the human WDR26-CTLH E3 ligase regulates prodrug metabolism. Mol Cell 84, 1948–1963 e1911. 10.1016/j.molcel.2024.04.014.

42. Barbulescu, P., Chana, C.K., Wong, M.K., Ben Makhlouf, I., Bruce, J.P., Feng, Y., Keszei, A.F.A., Wong, C., Mohamad-Ramshan, R., McGary, L.C., et al. (2024). FAM72A degrades UNG2 through the GID/CTLH complex to promote mutagenic repair during antibody maturation. Nat Commun 15, 7541. 10.1038/s41467-024-52009-x.

43. Grant, D.W., Tan, S., Ramage, D.E., Li, M.Z., Di, Y., Tchasovnikarova, I.A., Weekes, M.P., Elledge, S.J., Warren, A.J., and Timms, R.T. (2026). Proteome-wide C-degron activity profiling connects conditional regulation of the CTLH E3 ligase complex to ribosome biogenesis. bioRxiv. 10.64898/2026.01.14.698769.

44. Jeong, D.E., Lee, H.S., Ku, B., Kim, C.H., Kim, S.J., and Shin, H.C. (2023). Insights into the recognition mechanism in the UBR box of UBR4 for its specific substrates. Commun Biol 6, 1214. 10.1038/s42003-023-05602-7.

45. Hehl, L.A., Horn-Ghetko, D., Prabu, J.R., Vollrath, R., Vu, D.T., Perez Berrocal, D.A., Mulder, M.P.C., van der Heden van Noort, G.J., and Schulman, B.A. (2024). Structural snapshots along K48-linked ubiquitin chain formation by the HECT E3 UBR5. Nat Chem Biol 20, 190–200. 10.1038/s41589-023-01414-2.

46. Kim, H.K., Kim, R.R., Oh, J.H., Cho, H., Varshavsky, A., and Hwang, C.S. (2014). The N-terminal methionine of cellular proteins as a degradation signal. Cell 156, 158–169. 10.1016/j.cell.2013.11.031.

47. Timms, R.T., Zhang, Z., Rhee, D.Y., Harper, J.W., Koren, I., and Elledge, S.J. (2019). A glycine-specific N-degron pathway mediates the quality control of protein N-myristoylation. Science 365. 10.1126/science.aaw4912.

48. Grabarczyk, D.B., Ehrmann, J.F., Murphy, P., Yang, W.S., Kurzbauer, R., Bell, L.E., Deszcz, L., Neuhold, J., Schleiffer, A., Shulkina, A., et al. (2025). Architecture of the UBR4 complex, a giant E4 ligase central to eukaryotic protein quality control. Science 389, 909–914. 10.1126/science.adv9309.

49. Yang, Z., Haakonsen, D.L., Heider, M., Witus, S.R., Zelter, A., Beschauner, T., MacCoss, M.J., and Rape, M. (2025). Molecular basis of SIFI activity in the integrated stress response. Nature 643, 1117–1126. 10.1038/s41586-025-09074-z.

50. Heo, A.J., Kim, S.B., Ji, C.H., Han, D., Lee, S.J., Lee, S.H., Lee, M.J., Lee, J.S., Ciechanover, A., Kim, B.Y., and Kwon, Y.T. (2021). The N-terminal cysteine is a dual sensor of oxygen and oxidative stress. Proc Natl Acad Sci U S A 118. 10.1073/pnas.2107993118.

51. van Gen Hassend, P.M., and Schindelin, H. (2026). A structural code for assembly specificity in GID/CTLH-type E3 ligases. Elife 15. 10.7554/eLife.110152.

52. Sherpa, D., Mueller, J., Karayel, O., Xu, P., Yao, Y., Chrustowicz, J., Gottemukkala, K.V., Baumann, C., Gross, A., Czarnecki, O., et al. (2022). Modular UBE2H-CTLH E2-E3 complexes regulate erythroid maturation. Elife 11. 10.7554/eLife.77937.

53. Abramson, J., Adler, J., Dunger, J., Evans, R., Green, T., Pritzel, A., Ronneberger, O., Willmore, L., Ballard, A.J., Bambrick, J., et al. (2024). Accurate structure prediction of biomolecular interactions with AlphaFold 3. Nature 630, 493–500. 10.1038/s41586-024-07487-w.

54. Forelli, N., Thome, T., Eaton, D.M., Schultz, K., Patel, J., Bowman, C.E., Kawakami, R., Jung, J.W., Kuznetsov, I.A., Li, K., et al. (2026). SGLT2 inhibitors activate pantothenate kinase in the human heart. Science 393, 895–902. 10.1126/science.aeh4856.

55. Koren, I., Timms, R.T., Kula, T., Xu, Q., Li, M.Z., and Elledge, S.J. (2018). The Eukaryotic Proteome Is Shaped by E3 Ubiquitin Ligases Targeting C-Terminal Degrons. Cell 173, 1622–1635 e1614. 10.1016/j.cell.2018.04.028.

56. Carrillo Roas, S., Yagita, Y., Murphy, P., Kurzbauer, R., Clausen, T., Zavodszky, E., and Hegde, R.S. (2025). Convergence of orphan quality control pathways at a ubiquitin chain-elongating ligase. Mol Cell 85, 815–828 e810. 10.1016/j.molcel.2025.01.002.

57. Ji, C.H., Kim, H.Y., Heo, A.J., Lee, S.H., Lee, M.J., Kim, S.B., Srinivasrao, G., Mun, S.R., Cha-Molstad, H., Ciechanover, A., et al. (2019). The N-Degron Pathway Mediates ER-phagy. Mol Cell 75, 1058–1072 e1059. 10.1016/j.molcel.2019.06.028.

58. Pan, M., Zheng, Q., Wang, T., Liang, L., Mao, J., Zuo, C., Ding, R., Ai, H., Xie, Y., Si, D., et al. (2021). Structural insights into Ubr1-mediated N-degron polyubiquitination. Nature 600, 334–338. 10.1038/s41586-021-04097-8.

59. Matta-Camacho, E., Kozlov, G., Li, F.F., and Gehring, K. (2010). Structural basis of substrate recognition and specificity in the N-end rule pathway. Nat Struct Mol Biol 17, 1182–1187. 10.1038/nsmb.1894.

60. Mittl, P.R.E., and Beer, H.D. (2025). The B30.2/SPRY-Domain: A Versatile Binding Scaffold in Supramolecular Assemblies of Eukaryotes. Crystals 15. ARTN 281 10.3390/cryst15030281.

