## Supplementary figures for "Parallel Arg/N-degron recognition systems regulate coenzyme A biosynthesis"

### Supplemental Figures and Legends

#### Figure S1

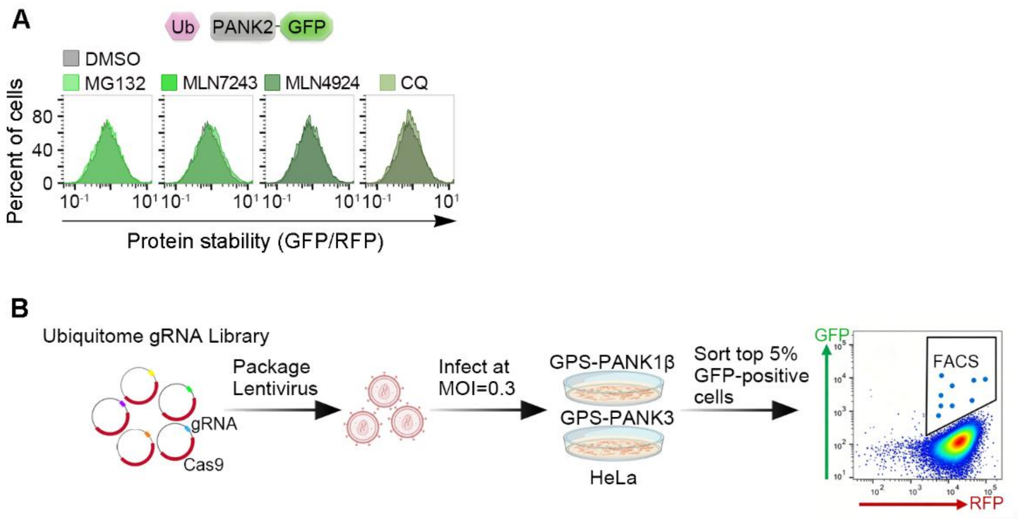


**Figure S1. PANK2 stability analysis using the global protein stability (GPS) reporter system and schematic overview of the CRISPR–Cas9 screening workflow. Related to Figure 1**

(A) Stable HeLa cell lines ectopically expressing GPS reporter for PANK2 were treated with 10 μM MG132, MLN7243, MLN4924 or CQ for 6h. Relative stability was quantified by flow cytometry based on the GFP/RFP fluorescence ratio.

(B) Schematic overview of the CRISPR-Cas9 screening workflow using GPS reporters and high-throughput sequencing to identify regulators of PANK1β or PANK3 protein stability.

#### Figure S2

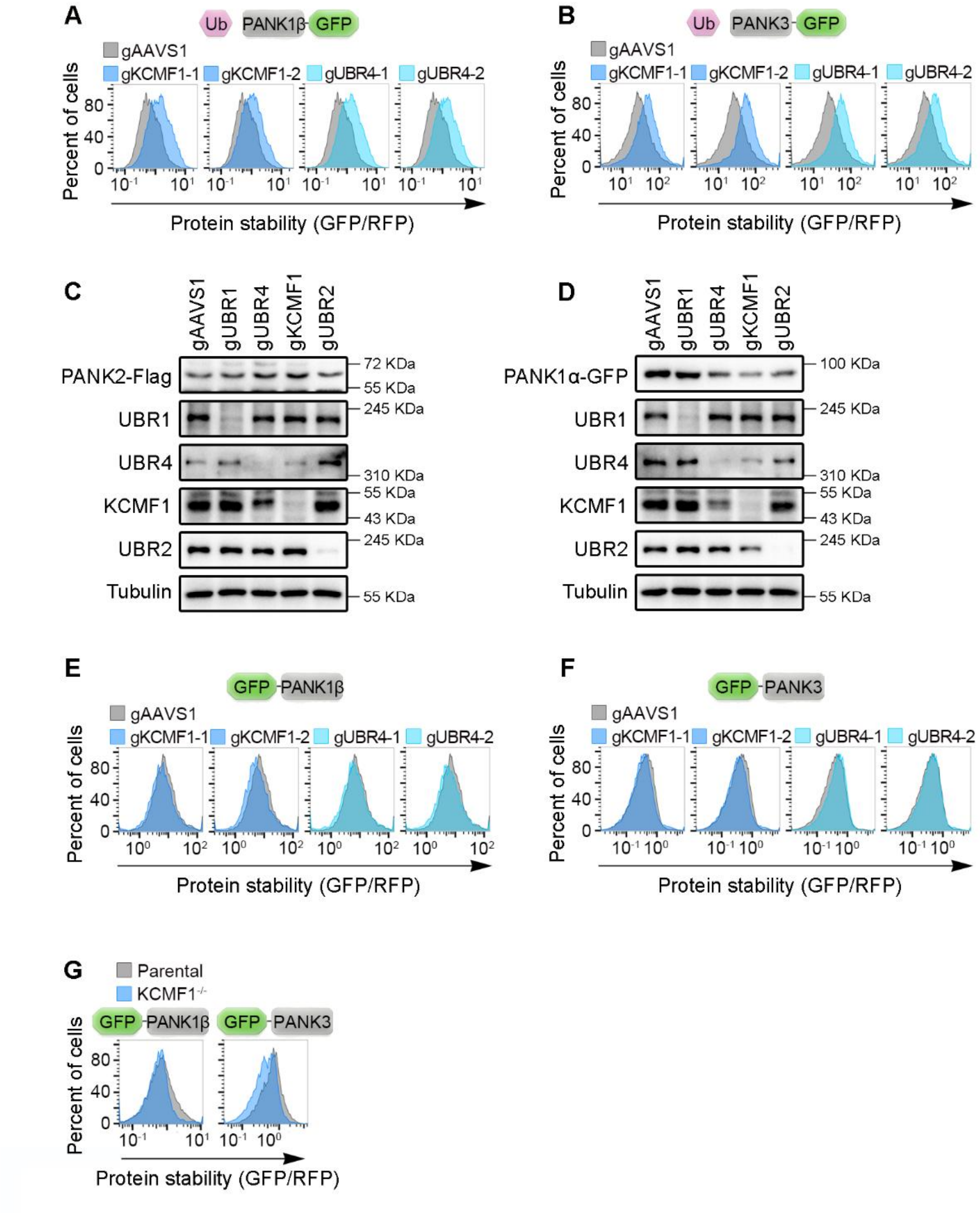


**Figure S2. UBR4–KCMF1 selectively promotes degradation of PANK1β and PANK3 through recognition of an N-terminal MK motif. Related to Figure 2**

(A) Stable HEK293T cells transduced with control gRNA or gRNA targeting KCMF1 or UBR4, as indicated, were transfected with PANK1β-GFP reporter, and reporter stability was assessed by flow cytometry.

(B) Stable HEK293T cells transduced with control gRNA or gRNA targeting KCMF1 or UBR4, as indicated, were transfected with PANK3-GFP reporter, and reporter stability was assessed by flow cytometry.

(C) Stable HEK293T cells ectopically expressing PANK2-Flag were transduced with control gRNA or gRNA targeting UBR1, UBR4, KCMF1 or UBR2, as indicated. Cell lysates were analyzed by immunoblotting with the indicated antibodies.

(D) Stable HEK293T cells ectopically expressing PANK1α-GFP were transduced with control gRNA or gRNA targeting UBR1, UBR4, KCMF1 or UBR2, as indicated. Cell lysates were analyzed by immunoblotting with the indicated antibodies.

(E) Stable HeLa cells transduced with control gRNA or gRNA targeting KCMF1 or UBR4, as indicated, were transfected with GFP-PANK1β reporter, and reporter stability was assessed by flow cytometry.

(F) Stable HEK293T cells transduced with control gRNA or gRNA targeting KCMF1 or UBR4, as indicated, were transfected with GFP-PANK3 reporter, and reporter stability was assessed by flow cytometry.

(G) WT and KCMF1$^{-/-}$ clonal HeLa cells were transfected with GFP-PANK1β or GFP-PANK3 reporters and analyzed by flow cytometry.

**Figure S3**

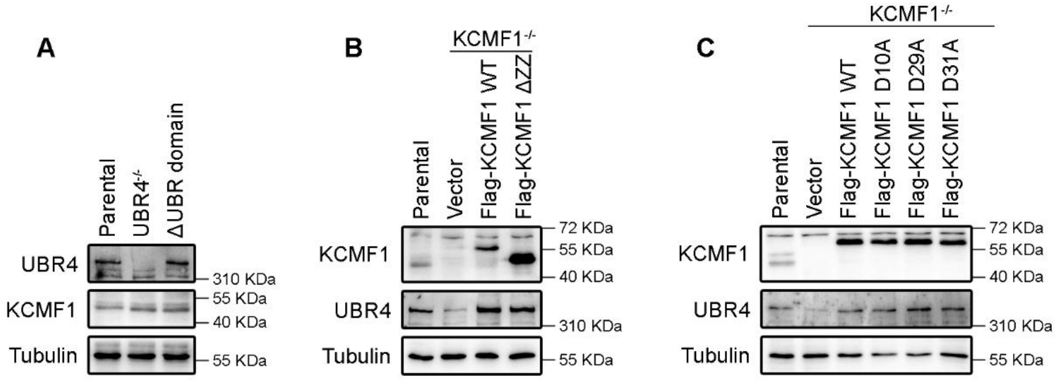


**Figure S3. Immunoblot validation of knockout and reconstituted cell lines. Related to Figure 3**

(A) Cell lysates from WT, UBR4$^{-/-}$, and UBR4 UBR domain-excised HEK293T cells were analyzed by immunoblotting with the indicated antibodies.

(B) Cell lysates from WT, KCMF1$^{-/-}$, and KCMF1$^{-/-}$ HEK293T cells re-expressing either WT KCMF1 or the ΔZZ mutant were analyzed by immunoblotting with the indicated antibodies.

(C) Cell lysates from WT, KCMF1$^{-/-}$, and KCMF1$^{-/-}$ HEK293T cells re-expressing WT KCMF1 or ZZ domain point mutants were analyzed by immunoblotting with the indicated antibodies.

**Figure S4**

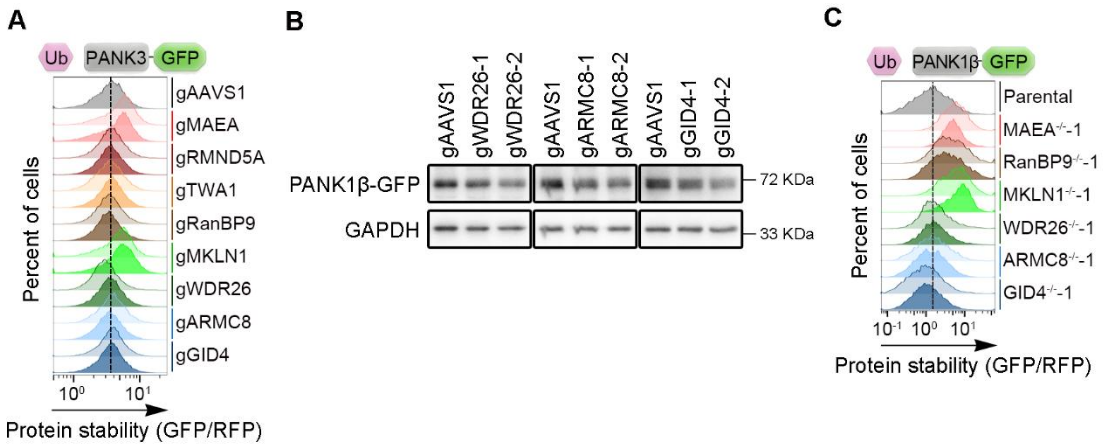


**Figure S4. The MKLN1-CTLH promotes PANK1α and PANK3 degradation. Related to Figure 4**

(A) WT and single-clone HEK293T cells deficient in the indicated CTLH complex components were transfected with the PANK3-GFP reporter and analyzed by flow cytometry.

(B) Stable HeLa cells ectopically expressing PANK1β-GFP were transduced with control gRNA or gRNA targeting WDR26, ARMC8 or GID4, as indicated. Cell lysates were analyzed by immunoblotting with the indicated antibodies.

(C) WT and knockout HeLa cells lacking the indicated individual CTLH complex components were transfected with PANK1β-GFP reporter, as indicated, and analyzed by flow cytometry.

**Figure S5**

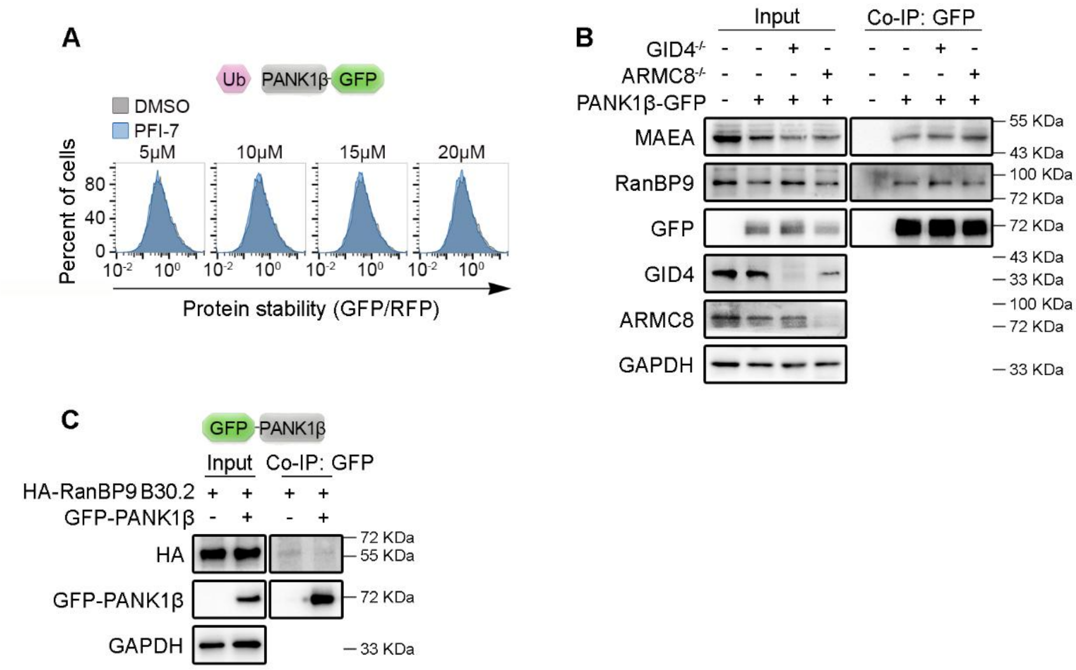


**Figure S5. The RanBP9 B30.2 domain associates with PANK1β. Related to Figure 5**

(A) Stable HEK293T cells ectopically expressing PANK1β-GFP reporter were treated with indicated concentrations of PFI-7 for 24 h and analyzed by flow cytometry.

(B) Parental HEK293T, GID4$^{-/-}$, or ARMC8$^{-/-}$ cells transiently expressing vector control or PANK1β-GFP were subjected to anti-GFP immunoprecipitation to examine interactions with endogenous MAEA and RanBP9. Input and immunoprecipitated samples were analyzed by immunoblotting with the indicated antibodies.

(C) HEK293T cells transiently co-expressing HA-tagged RanBP9 B30.2 domain (residues 1-359) together with an empty vector or GFP-PANK1β were subjected to anti-GFP co-IP. Input and co-IP samples were analyzed by immunoblotting with the indicated antibodies.

**Figure S6**

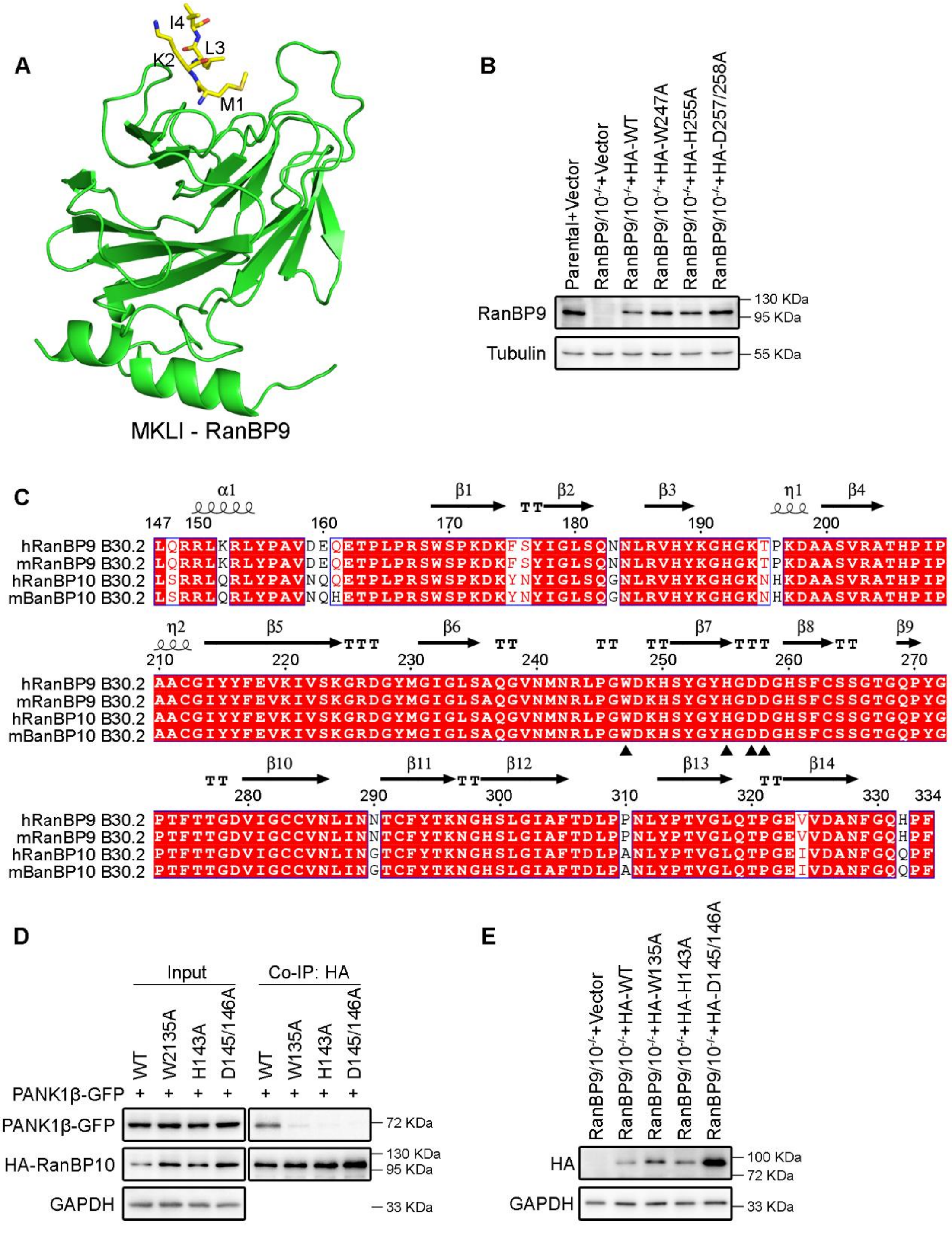


**Figure S6. Conserved B30.2 domains of RanBP9 and RanBP10 mediate PANK1β recognition. Related to Figure 6.**

(A) Overall structure of RanBP9 B30.2 domain in complex with the N-terminal MKLI motif of PANK1β. The MKLI peptide is shown as yellow sticks, and RanBP9 B30.2 domain is colored green.

(B) Cell lysates from WT, $RanBP9^{-/-}RanBP10^{-/-}$ and $RanBP9^{-/-}RanBP10^{-/-}$ HeLa cells re-expressing WT RanBP9 or the indicated B30.2 domain mutants were analyzed by immunoblotting with the indicated antibodies.

(C) Sequence alignment of B30.2 domains of human RanBP9 (Uniprot ID: Q96S59), mouse RanBP9 (Uniprot ID: P69566), human RanBP10 (Uniprot ID: Q6VN20) and mouse RanBP10 (Uniprot ID: Q6VN19).

(D) HEK293T cells transiently co-expressing HA-tagged WT or B30.2 domain mutant RanBP10 together with PANK1β-GFP were subjected to anti-HA co-IP. Input and co-IP samples were analyzed by immunoblotting with the indicated antibodies.

(E) Cell lysates from WT, RanBP9$^{-/-}$RanBP10$^{-/-}$ and RanBP9$^{-/-}$RanBP10$^{-/-}$ HeLa cells re-expressing WT RanBP10 or the indicated B30.2 domain mutants were analyzed by immunoblotting with the indicated antibodies.

**Figure S7**

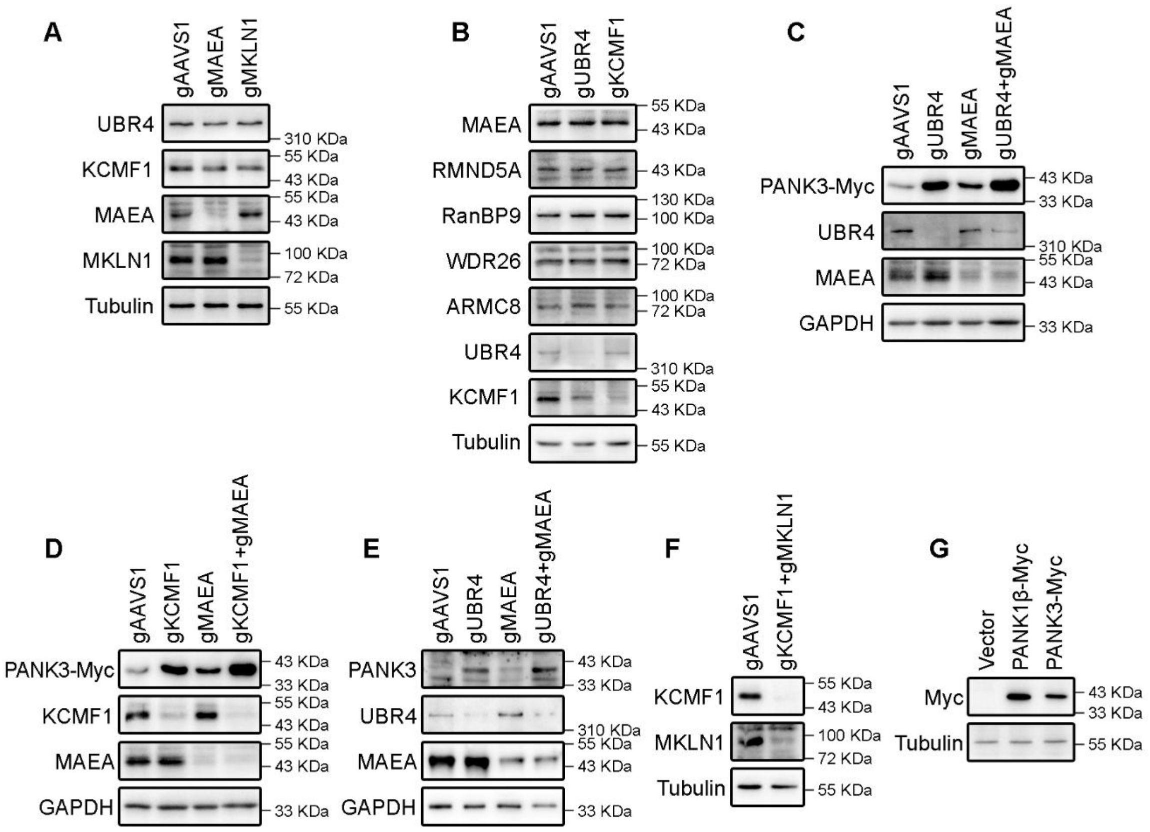


**Figure S7. UBR4–KCMF1 and MKLN1–CTLH independently mediate to PANK turnover. Related to Figure 7**

(A) Cell lysates from HEK293T cells transduced with control gRNA or gRNA targeting MAEA or MKLN1, as indicated, were analyzed by immunoblotting with the indicated antibodies.

(B) Cell lysates from HEK293T cells transduced with control gRNA or gRNA targeting UBR4 or KCMF1, as indicated, were analyzed by immunoblotting with the indicated antibodies.

(C) Stable HeLa cells ectopically expressing PANK3-Myc were transduced with control gRNA or gRNA targeting UBR4, MAEA or both genes simultaneously, as indicated. Cell lysates were analyzed by immunoblotting with the indicated antibodies.

(D) Stable HeLa cells ectopically expressing PANK3-Myc were transduced with control gRNA or gRNA targeting KCMF1, MAEA or both genes simultaneously, as indicated. Cell lysates were analyzed by immunoblotting with the indicated antibodies.

(E) HeLa cells were transduced with control gRNA or gRNA targeting UBR4, MAEA, or both genes simultaneously, as indicated. Cell lysates were analyzed by immunoblotting with the indicated antibodies.

(F) Cell lysates from WT and KCMF1/MKLN1 double-knockout HEK293T cells were analyzed by immunoblotting with the indicated antibodies.

(G) Cell lysates from HEK293T cells ectopically expressing an empty vector, PANK1β-Myc or PANK3-Myc were analyzed by immunoblotting with the indicated antibodies.

**Figure S8**

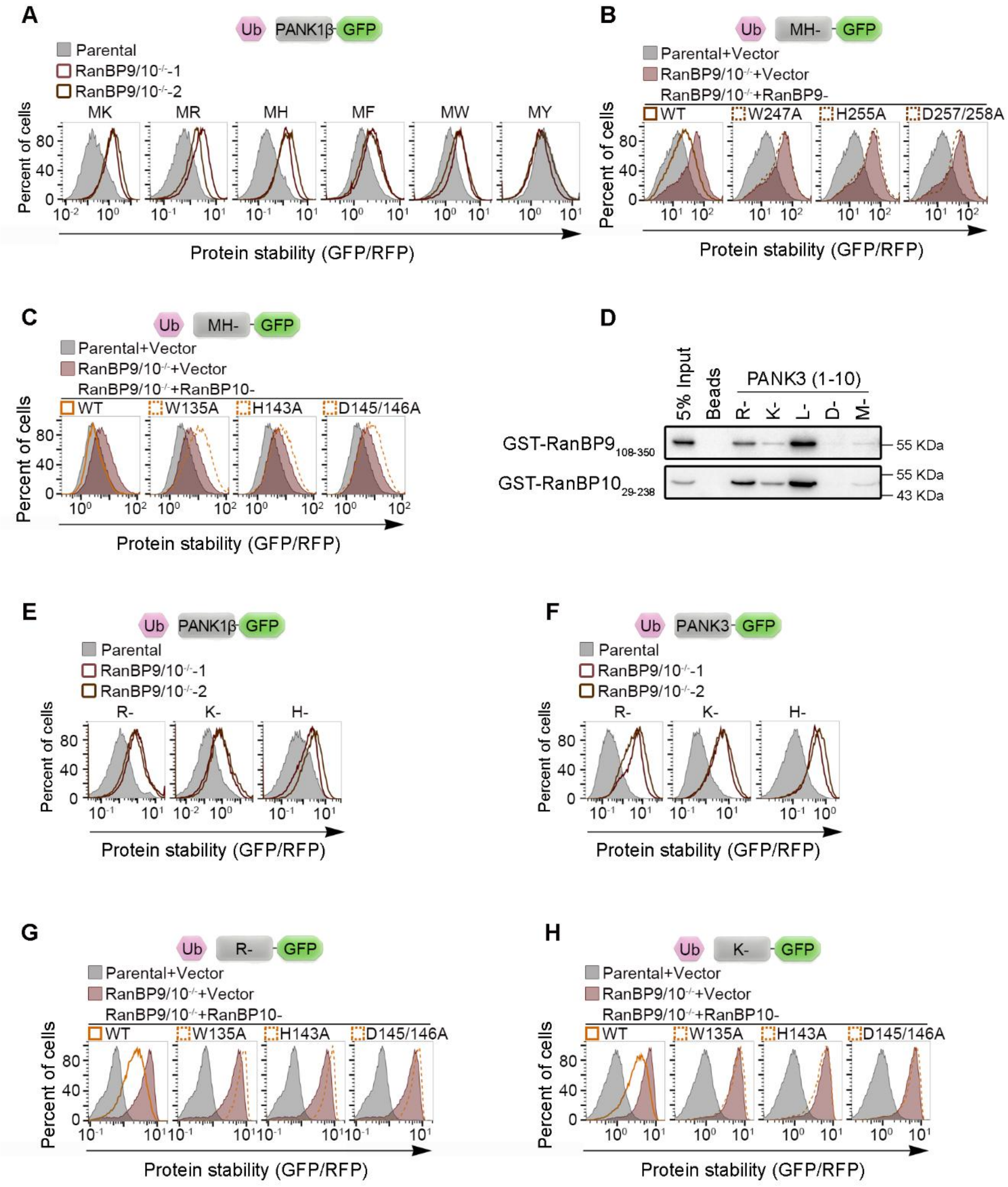


**Figure S8. RanBP9/RanBP10 recognize Arg/N-degrons. Related to Figure 8**

(A) WT and RanBP9$^{-/-}$RanBP10$^{-/-}$ HEK293T cells were transfected with PANK1β reporter variants bearing the indicated N-terminal MK-, MR-, MH-, MF-, MW- or MY-initiating sequences and analyzed by flow cytometry. Two independent knockout clones were examined for each genotype.

(B) WT, RanBP9$^{-/-}$RanBP10$^{-/-}$ and RanBP9$^{-/-}$RanBP10$^{-/-}$ HeLa cells reconstituted with WT RanBP9 or the indicated B30.2 domain mutants were transfected with MH-initiating PANK1β-GFP variant and analyzed by flow cytometry.

(C) WT, RanBP9$^{-/-}$RanBP10$^{-/-}$ and RanBP9$^{-/-}$RanBP10$^{-/-}$ HeLa cells reconstituted with WT RanBP10 or the indicated B30.2 domain mutants were transfected with MH-initiating PANK1β-GFP variant and analyzed by flow cytometry.

(D) Peptide pull-down assay using the indicated biotinylated PANK3 N-terminal peptides carrying substitutions at the first residue, incubated with GST-fused RanBP9 or RanBP10 B30.2 domains.

(E) WT and RanBP9$^{-/-}$RanBP10$^{-/-}$ HEK293T cells were transfected with PANK1β reporter variants bearing the indicated N-terminal R-, K- or H- initiating sequences and analyzed by flow cytometry. Two independent knockout clones were examined for each genotype.

(F) WT and RanBP9$^{-/-}$RanBP10$^{-/-}$ HEK293T cells were transfected with PANK3 reporter variants bearing the indicated N-terminal R-, K- or H- initiating sequences and analyzed by flow cytometry. Two independent knockout clones were examined for each genotype.

(G) WT, RanBP9$^{-/-}$RanBP10$^{-/-}$ and RanBP9$^{-/-}$RanBP10$^{-/-}$ HeLa cells reconstituted with WT RanBP10 or the indicated B30.2 domain mutants were transfected with the R-initiating PANK3-GFP variant reporter and analyzed by flow cytometry.

(H) WT, RanBP9$^{-/-}$RanBP10$^{-/-}$ and RanBP9$^{-/-}$RanBP10$^{-/-}$ HeLa cells reconstituted with WT RanBP10 or the indicated B30.2 domain mutants were transfected with the K-starting PANK3-GFP variant reporter and analyzed by flow cytometry.
